# ZMYND11 Functions in Bimodal Regulation of Latent Genes and Brain-like Splicing to Safeguard Corticogenesis

**DOI:** 10.1101/2024.10.15.618524

**Authors:** Xuyao Chang, Wenqi Li, Satoshi Matsui, Cindy Huynh, Gustav Y. Cederquist, Lorenz Studer, Makiko Iwafuchi, Amelle Shillington, Constantinos Chronis, Jason Tchieu

## Abstract

Despite the litany of pathogenic variants linked to neurodevelopmental disorders (NDD) including autism (ASD) and intellectual disability^1,2^, our understanding of the underlying mechanisms caused by risk genes remain unclear. Here, we leveraged a human pluripotent stem cell model to uncover the neurodevelopmental consequences of mutations in *ZMYND11*, a newly implicated risk gene^3,4^. ZMYND11, known for its tumor suppressor function, encodes a histone-reader that recognizes sites of transcriptional elongation and acts as a co-repressor^5,6^. Our findings reveal that ZMYND11-deficient cortical neural stem cells showed upregulation of latent developmental pathways, impairing progenitor and neuron production. In addition to its role on histones, ZMYND11 controls a brain-specific isoform switch involving the splicing regulator RBFOX2. Extending our findings to other chromatin-related ASD risk factors revealed similar developmental pathway activation and splicing dysregulation, partially rescuable through ZMYND11’s regulatory functions.

## Main Text

Chromatin and epigenetic-related genes play a crucial role in shaping this developmental landscape and influencing future cell fate decisions^7–10^. Despite their broad importance, genomic studies have found that several chromatin-related genes are mutated in autism spectrum disorders (ASD), suggesting their potential contribution to neurodevelopmental vulnerability^1,2^. *De novo* pathogenic variants in ZMYND11, a newly identified neurodevelopmental disorder (NDD), is associated with intellectual disability, epilepsy, and autistic traits^3,4^. ZMYND11 is a chromatin reader that targets transcriptional elongation sites by recognizing and binding to modified histones marked by H3K36me3^5,6^. Once bound, ZMYND11 acts as a transcriptional repressor by inhibiting transcription, which has been observed in cancer models. However, its role in cortical brain development is unexplored.

Intriguingly, elevated expression of ZMYND11 in breast cancer has been linked to improved survival, underscoring its crucial role as a tumor suppressor^6,11^. However, it remains unknown whether similar repressive mechanisms, when disrupted, confer risk for NDD. Our study investigates the transcriptional repressor role of ZMYND11 during human cortical brain development, utilizing a pluripotent stem cell derived model of cortical neurogenesis^12–15^. We identify ZMYND11 as a key regulator at the intersection of epigenetic control and alternative splicing, mediating a critical switch between brain-specific and non-brain mRNA isoforms in cortical neural stem cells (NSCs). Notably, this dysregulation in tissue specific mRNA isoforms is found across multiple high-confidence chromatin-related ASD risk factors, suggesting broader implications for the etiology of both ASD and NDD.

### ZMYND11 is critical for the differentiation of radial glial neural stem cells (NSCs) into intermediate progenitor cells (IPCs), a process impaired in a subset of autism risk genes

Pathogenic variants within the coding region of *ZMYND11* predominantly cause frameshift mutations, leading to haploinsufficiency^3,4^ (**Extended Data Fig. 1a**). *ZMYND11* expression is present throughout fetal brain development, suggesting *de novo* mutations may affect early cortical brain development. To determine its role in this process, we engineered loss-of-function and heterozygous/patient-like mutations in human embryonic stem cells (hESCs) to investigate its function in disease-relevant cell types within an isogenic background (**Extended Data Fig. 1b**). We validated mutation status and confirmed altered protein expression (**Extended Data Fig. 1c-e**). These novel lines were karyotypically normal (**Extended Data Fig. 1f**), did not reveal noteworthy changes in the expression of pluripotency markers such as NANOG and SOX2 (**Extended Data Fig. 1g**), nor were there significant differences in the cell proliferation (**Extended Data Fig. 1h**).

We next asked whether there were any cellular phenotypes associated with ZMYND11 deficiency during the differentiation into cortical neural stem cells (NSCs). Many ASD risk genes fall into classes where cortical NSCs delay (Class I) or accelerate (Class II) cortical development^13^. To determine which class *ZMYND11* is more associated with, we generated cortical organoids using a guided differentiation approach (**Fig. 1a**). We were able to obtain cortical organoids that represented the anterior forebrain marked by FOXG1 expression, that also expressed intermediate progenitor (IPC) marker, TBR2, indicative of cells undergoing corticogenesis (**Fig. 1b**). We found a dramatic decrease in FOXG1 and TBR2 expression in the loss of function line suggesting that ZMYND11 could play a role in neural patterning (**Fig. 1c**).

**Fig. 1.**
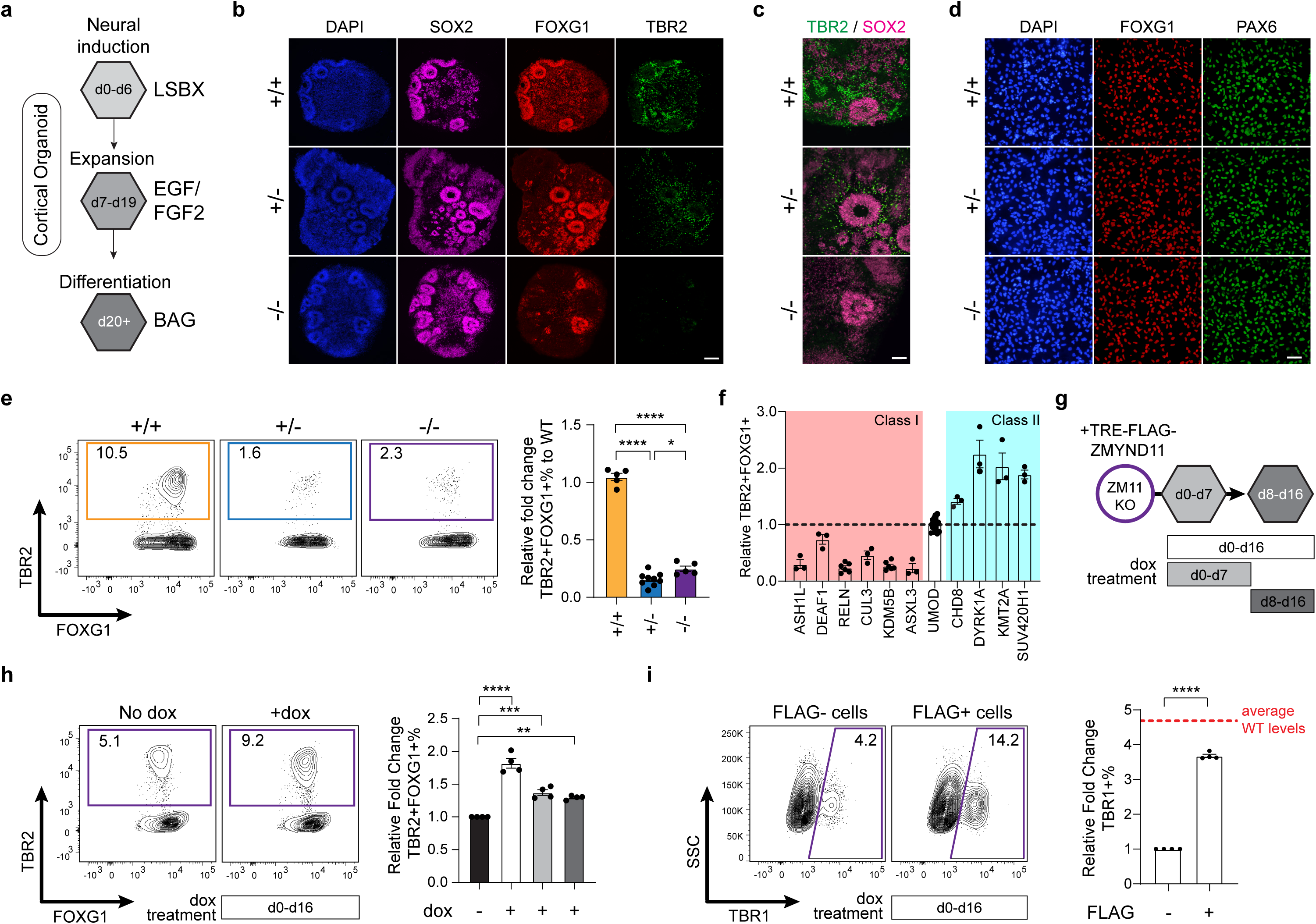
Engineered human stem cell model of ZMYND11 NDD reveals a dramatic decrease in the production of intermediate progenitors (IPCs) upon cortical differentiation. **a.** Schematic outline of three-dimensional cortical organoid differentiation protocol. LSBX: LDN193189 (BMPi), SB431542 (TGF-βi), XAV939 (WNTi). BAG: BDNF, Ascorbic Acid, GDNF. **b-c**. Immunofluorescence staining for SOX2, FOXG1 and TBR2 on WT and ZMYND11 deficient cortical organoid section at d30 (**b**). Zoomed-in view (**c**). Scale bars, 200μm in **b**, 100μm in **c**. **d**. Immunofluorescence staining for FOXG1 and PAX6 on WT and ZMYND11 deficient monolayer culture at d18. Scale bars, 50μm. **e**. Intracellular flow cytometry analysis for TBR2 and FOXG1 on WT and ZMYND11 deficient monolayer culture at d16 (left) with quantification relative to WT (right, mean±SEM, n=5 differentiations for +/+ and -/-, n=9 differentiations using two clones for +/-). **f**. Quantification of intracellular flow cytometry analysis for TBR2 and FOXG1 on Class I mutants (n=3 differentiations for *ASH1L, DEAF1, CUL3, ASXL3*; n=6 differentiations using two clones for *RELN, KDM5B*) and Class II mutants (n=3 differentiations for *CHD8, DYRK1A, KMT2A, SUV420H1*) relative to isogenic control (*UMOD* edited lines, n=15 differentiations using three clones, mean±SEM) monolayer culture at d16. **g**. Schematic outline for overexpression of FLAG-ZMYND11 in ZMYND11 KO using dox-inducible system (200ng/ml) either throughout the differentiation (d0-d16) or during half of the differentiation (d0-d7 or d8-d16). **h**. Intracellular flow cytometry analysis for TBR2 and FOXG1 on no doxycycline and dox-treated monolayer culture at d16 (left) with quantification relative to no dox treated group (right, mean±SEM, n=4 differentiations). **i**. Intracellular flow cytometry analysis for TBR1 on FLAG- and FLAG+ populations on d0-d16 dox-treated monolayer culture at d16 (left) with quantification relative to FLAG-population (right, mean±SEM, n=4 differentiations). Red dotted line marks average WT levels calculated by TBR1% fold decrease in ZMYND11 KO relative to WT. Statistics in **e**, **h**: one-way ANOVA followed by Tukey’s test. Statistics in **i**: unpaired two-tailed *t* test (two groups). \**P*<0.05; \*\**P*<0.01; \*\*\**P*<0.001; \*\*\*\**P*<0.0001. *P* values in **e**: +/- vs +/+: *P*<0.0001, -/- vs +/+: *P*<0.0001, -/- vs +/-: *P*=0.035; in **h**: d0-d16 dox vs no dox: *P*<0.0001, d0-d7 dox vs no dox: *P*=0.0003, d8-d16 dox vs no dox: *P*=0.0023; in i: FLAG+ vs FLAG-: *P*<0.0001.

To adopt a more quantitative approach and reduce the potential variability found within cortical organoid culture, we performed monolayer differentiations in conjunction with intracellular flow cytometry to measure changes in IPC generation. In contrast to cortical organoids, the directed differentiation of hESCs into prefrontal-like cortical NSCs (**Extended Data Fig. 2a, b**) was highly efficient in all lines (**Fig. 1d and Extended Data Fig. 2c, d**). From these populations, we observed a 4-7-fold decrease in IPC generation in *ZMYND11* mutants (**Fig. 1e**), a pattern similarly found in Class I risk genes (**Fig. 1f**). This reduction in IPCs correlates with an abnormal retention of cortical NSC identity, marked by expression of CD133 (**Extended Data Fig. 2e**), indicating a resistance to progress normally through cortical differentiation rather than undergoing spontaneous differentiation into neurons. To confirm this, we found that the generation of TBR1-positive deep layer neurons was also impaired to a similar magnitude (**Extended Data Fig. 2f**). Intriguingly, *ZMYND11* mutants did not display differences in IPC or neuron differentiation between the heterozygous and the knockout lines, suggesting that once ZMYND11 levels fall below a critical threshold, further reduction does not exacerbate the impact on cortical differentiation.

To test whether re-expression of ZMYND11 in the knockout line could rescue the IPC phenotype, we cloned ZMYND11 with an N-terminal 3X FLAG tag into a doxycycline-inducible vector and activated it during differentiation (**Fig. 1g**). We found that expression of FLAG-ZMYND11 throughout differentiation led to the highest yield of IPCs and subsequent neuron differentiation (∼1.8 increase, **Fig. 1h**) compared to expression during only half of differentiation (∼1.3 increase). Due to the silencing of the transgene in cortical NSCs, we examined transduced FLAG expressing cells and compared them to cells lacking FLAG (**Extended Data Fig. 2g**). We found that neuron differentiation in FLAG+ NSCs exhibited ∼4-fold increase, similar to the magnitude of decrease in ZMYND11 knockout (red dotted line, **Fig. 1i**). This suggests that the IPC deficient and differentiation phenotypes in *ZMYND11* mutants can be reversed by restoring expression.

### Elevated expression of developmental signaling transcripts impairs corticogenesis in ZMYND11 deficient cortical NSCs

To determine the underlying molecular changes associated with ZMYND11 deficiency, we performed bulk RNA sequencing due to our highly homogenous population of cortical NSCs as well as the starting hESCs. Principal component analysis revealed that cell types clustered together based on their cell identity, regardless of genotype differences but, the magnitude of differential gene expression was more pronounced within the cortical NSC population (mutants vs controls) (**Fig. 2a and Supplementary Table 1**). As expected, differential gene regulatory networks between hESCs and cortical NSCs converged on gene ontologies related to neural differentiation (**Extended Data Fig. 3a**). Gene expression changes between controls and ZMYND11 deficient cortical NSCs significantly overlapped (**Fig. 2b**) and since ZMYND11 is a known transcriptional repressor, we focused on genes differentially upregulated. ZMYND11 deficient NSCs displayed significant upregulation in gene networks associated with the BMP and WNT signaling pathways (**Fig. 2c and Extended Data Fig. 3b**) which may explain the enrichment for additional ontologies associated with non-nervous system developmental processes (**Extended Data Fig. 3c**). We validated the increase in BMP signaling in mutant lines by measuring levels of phosphorylated SMAD1/5/8 (**Fig. 2d**) and found this correlated with impaired neuron differentiation when we treated the cells with a γ-secretase inhibitor, DAPT (**Fig. 2e, f**).

**Fig. 2.**
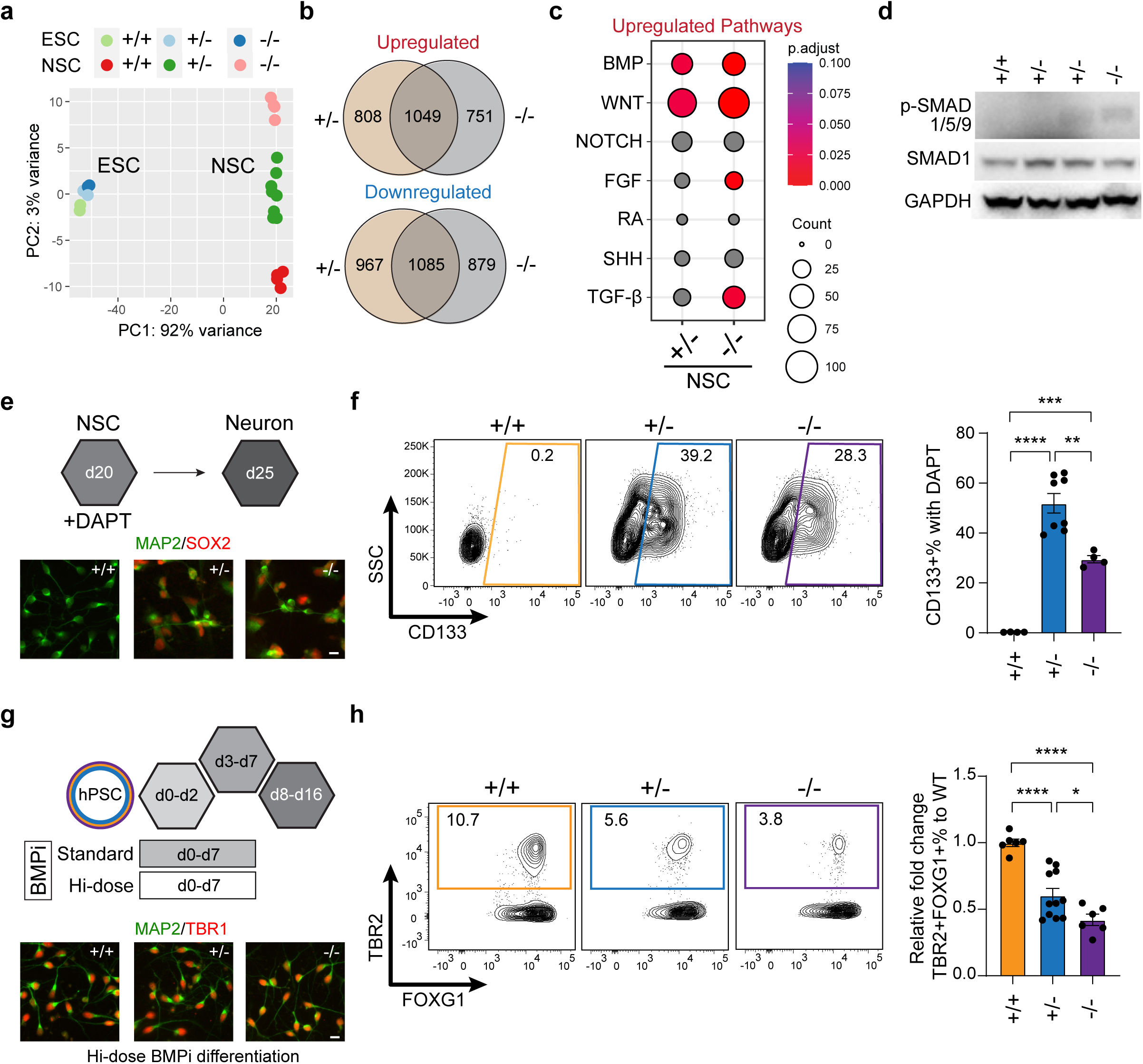
Transcriptomic analysis reveals elevated levels of BMP and WNT signaling pathways, when modulated, can partially rescue IPC production in ZMYND11 deficient cortical neural stem cells (NSCs) **a**. Principal component analysis of RNA-seq on WT and ZMYND11 deficient hESCs and NSCs (for hESCs, n=2 replicates for WT and -/-, n=2 clones for +/-; for NSCs, n=4 differentiations for +/+ and -/-; n=8 differentiations using two clones for +/-). **b.** Overlap of differentially expressed genes in NSCs on ZMYND11 +/- vs WT and ZMYND11 -/- vs WT NSCs (hypergeometric *P* value < 1e-100). **c.** Gene ontology analysis on upregulated genes in ZMYND11 deficient NSCs using biological processes focusing on growth factor signaling pathways. *P*.adjust values were calculated using Benjamini-Hochberg method. **d.** Western blot on p-SMAD1/5/8 and SMAD1 on ZMYND11 deficient NSCs. **e.** Scheme outlining neuron induction strategy from NSCs (upper) and immunofluorescence staining for MAP2 and SOX2 on WT and ZMYND11 deficient cultures after neuron induction (lower). Scale bars, 10μm. **f.** Flow cytometry analysis for CD133 on WT and ZMYND11 deficient neuron induction culture at d25 (left) with quantification (right, mean±SEM, n=4 differentiations for WT and -/-, n=4 differentiations using two clones for +/-). **g.** Scheme outlining high-dose BMP inhibition strategy (LDN193189: 500nM) during monolayer differentiation (upper) and immunofluorescence staining for MAP2 and TBR1 on WT and ZMYND11 deficient cultures after neuron induction (lower). Scale bars, 10μm. **h.** Intracellular flow cytometry analysis for TBR2 and FOXG1 on WT and ZMYND11 deficient monolayer culture after high-dose BMP inhibition at d16 (left) with quantification relative to WT (right, mean±SEM, n=6 differentiations for WT and -/-, n=11 differentiations using two clones for +/-). Statistics in **f**, **h**: one-way ANOVA followed by Tukey’s test. \**P*<0.05; \*\**P*<0.01; \*\*\**P*<0.001; \*\*\*\**P*<0.0001. *P* values in **f**: +/- vs +/+: *P*<0.0001, -/- vs +/+: *P*=0.0006, -/- vs +/-: *P*=0.0018; *P* values in **h**: +/- vs +/+: *P*<0.0001, -/- vs +/+: *P*<0.0001, -/- vs +/-: *P*=0.0194.

Due to these potent signaling molecules ability to alter cell fate, we modulated WNT and BMP signaling independently to retain cortical identity (**Extended Data Fig. 4a, b**). For the Hi-dose BMP inhibition, ZMYND11 deficient cortical NSCs were able to differentiate into deep layer excitatory neurons identified by TBR1- and VGLUT2-expression after DAPT treatment (>90%, **Fig. 2g and Extended Data Fig. 4c, d**). We next wondered whether ectopic BMP signaling could also impair IPC generation. To test this, we performed differentiations with the Hi-dose level of BMP inhibitor and found an increase in IPC generation from ZMYND11 deficient NSCs with no change in IPC numbers in control cells (**Fig. 2h**). Conversely, when we extended the duration of the WNT signaling inhibitor, XAV939, we surprisingly found no significant change in IPC generation (**Extended Data Fig. 4e, f**). Finally, to determine which lineages were enriched during spontaneous differentiation of *ZMYND11* mutants and isogenic controls, we performed an embryoid body formation assay. Within 10 days of differentiation, we observed a profound upregulation of mesenchymal rather than epithelial gene expression suggesting that gene regulatory networks associated with mesendodermal-like fate dominate which could factor in disrupting the generation of cortical organoids from knockout lines (**Extended Data Fig. 3d**). These data demonstrate that ZMYND11 deficiency elevates the expression of developmental signaling pathways that impair cortical NSCs to differentiate into IPCs and neurons. This impairment can be partially rescued by further inhibition of BMP, but not WNT, signaling pathways.

### ZMYND11 binding represses latent gene regulatory networks associated with BMP and WNT signaling pathways

ZMYND11 is known to binds sites of transcriptional elongation identified by histones modified with trimethylation of histone 3 (H3K36me3) to prevent transcription^6^. To determine whether these transcriptomic changes are directly caused by ZMYND11 deficiency, we performed CUT&RUN in cortical NSCs for ZMYND11 and H3K36me3. We found that ZMYND11 largely occupied promoter sites and positively correlated with H3K36me3 (**Fig. 3a, b**). To reduce uncertainty associated with the ZMYND11 and H3K36me3 antibodies and this assay, we performed CUT&RUN for ZMYND11 in haploinsufficient and loss of function lines and found a reduction in overall binding and peak height, respectively confirming specificity and used the loss of function line as our negative control (**Extended Data Fig. 5a, b**). In parallel, we screened a panel of published H3K36me3 antibodies to identify those effective in CUT&RUN and chose the one with the highest enrichment at genes with active transcription (**Extended Data Fig. 5c**). While our H3K36me3 genomic binding profiles showed enrichment along the gene body, the ZMYND11 binding profiles differed somewhat from previous studies potentially reflecting cell type-specific variations in recruitment. To confirm this, we generated a doxycycline-inducible FLAG-tagged ZMYND11 and assessed its binding pattern in cortical NSCs (**Extended Data Fig. 5d**). After 3 days of induction, we observed that peaks called by FLAG and ZMYND11 had ∼80% overlap; as expected, the FLAG antibody yielded significantly more peaks compared to the ZMYND11 antibody alone and reproduced our initial observation of enriched promoter occupancy. In cortical NSCs, ZMYND11 seems to preferentially bind promoter regions, which may differ from its binding patterns in other cell types studied. Therefore, we reanalyzed all publicly available ZMYND11 binding data and found that out of six studies^5,6,16–19^, four displayed promoter enrichment, suggesting that ZMYND11 binding profiles may be cell type specific (**Extended Data Fig. 5e**).

**Fig. 3.**
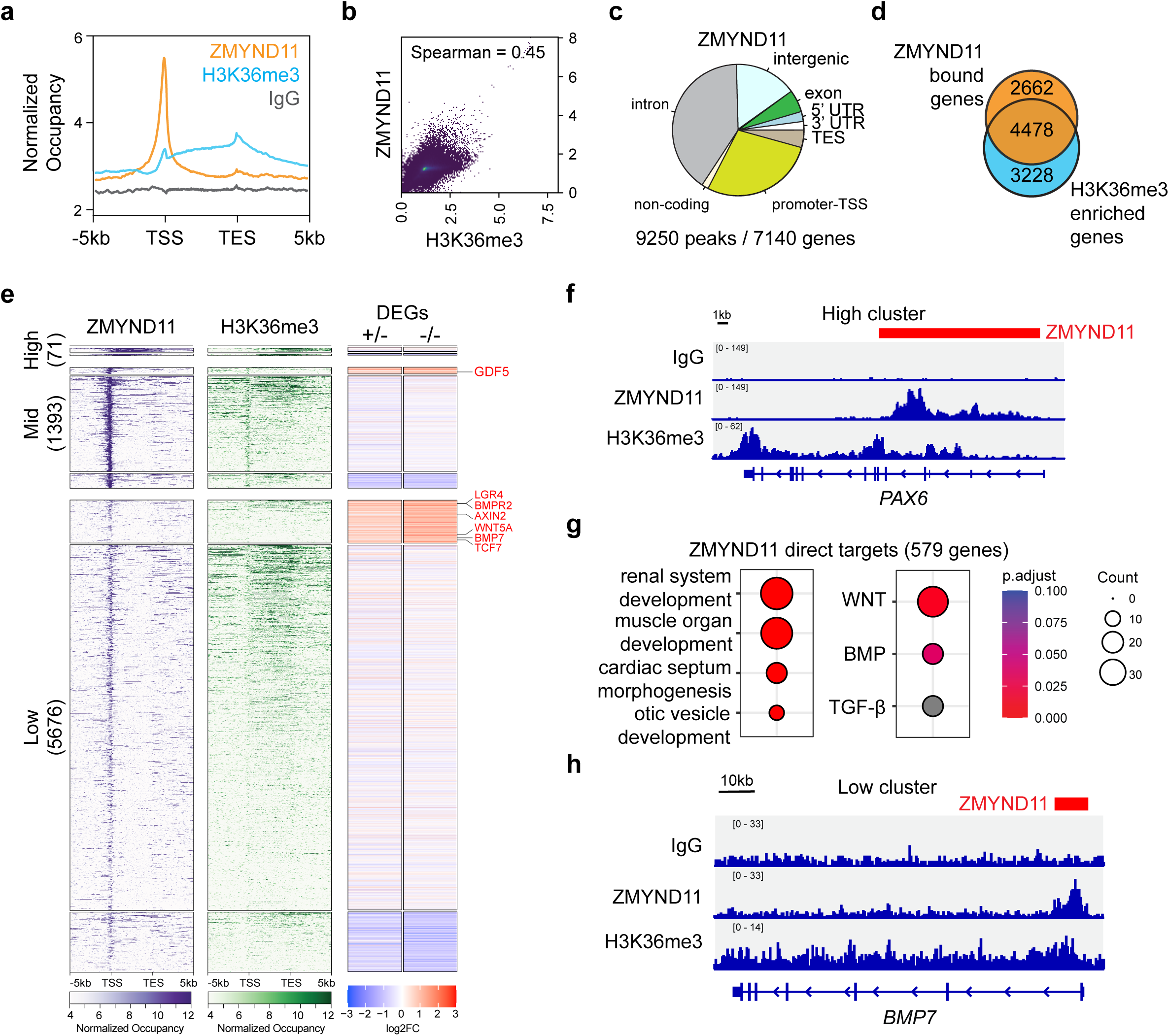
ZMYND11 directly binds and transcriptionally represses BMP and WNT signaling-related genes. **a.** Normalized ZMYND11 binding and H3K36me3 enrichment in WT cortical NSCs from transcription start sites (TSS) to transcription end sites (TES) ± 5kb (n=2 replicates). **b.** Correlation plot of ZMYND11 binding and H3K36me3 enrichment in WT NSCs (correlation coefficient calculated using Spearman method). **c.** Genomic features distribution, number of called peaks and bound genes of ZMYND11 binding in WT NSCs. **d.** Overlap of ZMYND11 bound genes with H3K36me3 enriched genes (hypergeometric *P* value < 1e-100). **e.** k-means clustering of ZMYND11 binding for ZMYND11 bound genes (High group=71 genes, Mid group=1393 genes, Low group=5676 genes), related H3K36me3 enrichment and heatmap plotting gene expression log2FC (fold change) by RNA-seq. Genes within each group were further separated according to gene expression changes (upregulated, no change, downregulated). Selected genes related with BMP and WNT signaling pathways are marked in red. **f.** Representative track (*PAX6*) of ZMYND11 and H3K36me3 from high ZMYND11 binding cluster. Red marks called ZMYND11 peak. **g.** Gene ontology analysis on ZMYND11-bound genes that were upregulated in KO (579 genes, ZMYND11 direct targets) using biological processes. *P*.adjust values were calculated using Benjamini-Hochberg method. **h.** Representative track (*BMP7*) of ZMYND11 and H3K36me3 that were related to BMP & WNT from low ZMYND11 binding cluster. Red marks called ZMYND11 peak.

ZMYND11 and H3K36me3 bind to 7,140 and 7,706 genes in cortical NSCs (**Fig. 3c, Extended Data Fig. 6a and Supplementary Table 2**), respectively with an overlap of 4,478 genes (**Fig. 3d**). Combining our transcriptomic and binding data identified three major clusters that encompass ZMYND11 binding patterns. The “High” cluster binds across the gene body, the “Mid” cluster is strongly enriched at the promoter, and the “Low” cluster displays weak promoter binding (**Fig. 3e**). Several critical genes associated with cortical NSC identity such as PAX6 are in High cluster (**Fig. 3f**). Surprisingly, by correlating binding with expression we found that ZMYND11 deficiency has little effect on global gene expression (**Extended Data Fig. 6b**). Interestingly, of the three clusters analyzed, we found that genes with weak ZMYND11 binding at the promoter (Low cluster) had significant changes in gene expression and are associated at sites with less H3K36me3 co-binding (**Extended Data Fig. 6c**). These ZMYND11 bound direct targets were enriched in non-nervous system developmental programs and signaling pathways associated with WNT and BMP signaling (**Fig. 3g, h**). Genes related to extracellular matrix and mesenchyme development, which were also differentially regulated, were not bound suggesting that these changes are caused by an indirect mode of dysregulation (**Extended Data Fig. 6d**). These genomic data analyses suggest that ZMYND11 directly regulates genes associated with latent, non-nervous system development and related pathways at promoters in cortical NSCs.

### ZMYND11 deficiency correlates with an alternative splicing switch from brain to non-brain isoform expression

In addition to the role ZMYND11 plays on chromatin, we next focused on the significant alternative splicing changes observed within cortical NSCs. Knockdown studies in cancer lines have shown that dysregulation of ZMYND11 increases differential alternative splicing (DAS) events that may contribute to tumorigenesis^5,11^. To determine whether ZMYND11 deficient hESCs and cortical NSCs exhibit DAS, we performed splicing analysis using our transcriptomic datasets and found significant differential exon usage in ZMYND11 deficient cortical NSCs compared to controls and lower DAS events in hESCs (**Fig. 4a and Extended Data Fig. 7a**). We focused on cortical NSCs and the DAS events predominantly involved spliced exon events. We next plotted principal components of exon usage, revealing a distinct separation of exon usage compared to controls (**Fig. 4b**). To minimize false positives from sequence alignment and statistical prediction algorithms, we performed additional analysis using two additional pipelines and considered only the intersecting events (**Extended Data Fig. 7b**). This approach identified 991 and 590 DAS events in heterozygous and homozygous mutations, respectively (**Fig. 4c, Extended Data Fig. 7c, d and Supplementary Table 3**), exhibiting a significant overlap (**Extended Data Fig. 7e**). Interestingly, the same approach found only fewer overlapped DAS events in mutant hESCs **(Extended Data Fig. 7f**). Upon examining potential shared properties of the spliced isoforms in cortical NSCs, we found a significant correlation with the inclusion or exclusion of small exons (**Extended Data Fig. 7g**), which has been described in prior work on autism^20,21^.

**Fig. 4.**
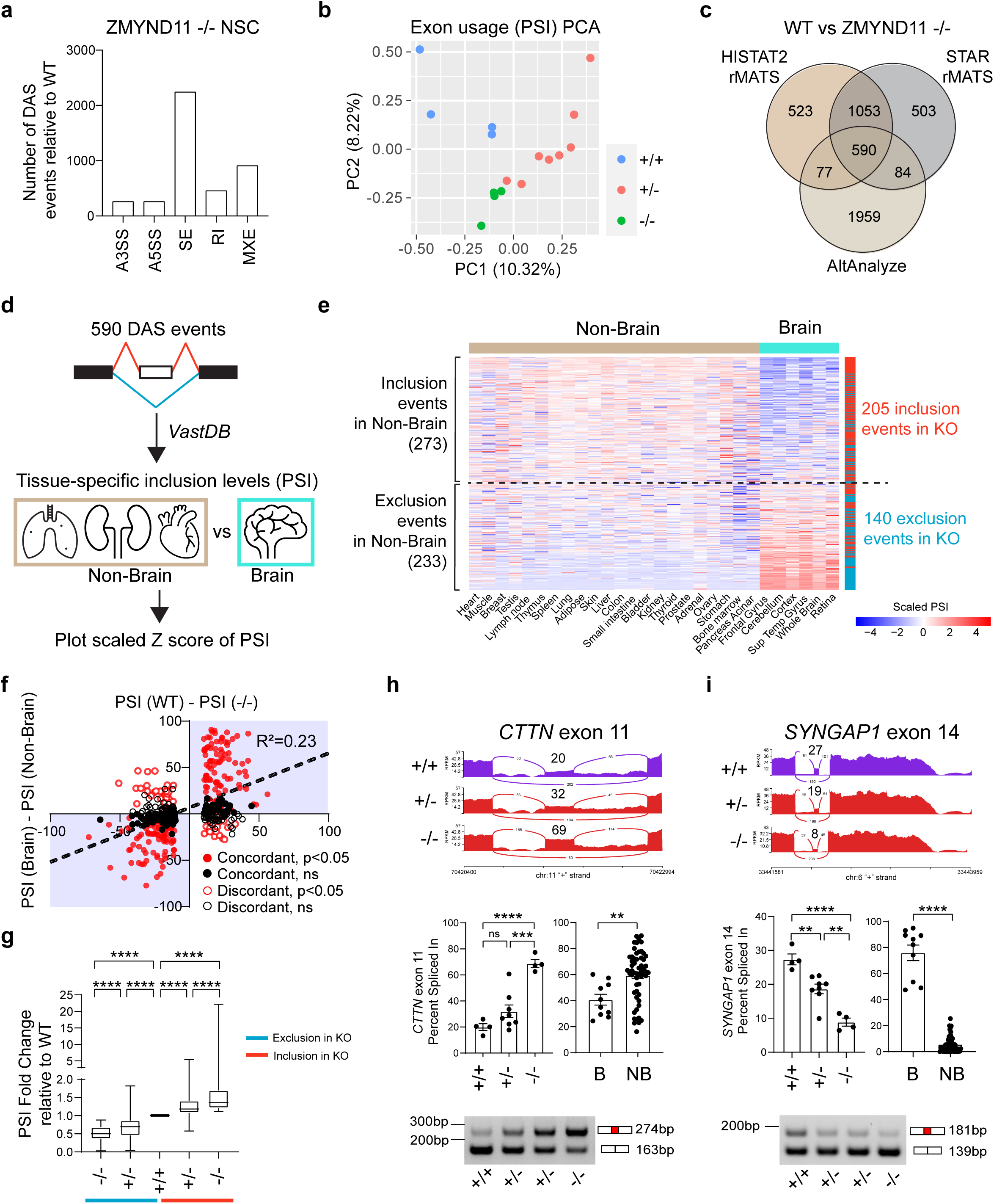
ZMYND11 promotes a brain-specific mRNA isoform switch. **a.** Number of significantly changed differential alternative splicing (DAS) events in ZMYND11 -/- cortical NSCs using rMATS prediction and HISAT2 as alignment software. A3SS: Alternative 3’ splice site. A5SS: Alternative 5’ splice site. SE: Spliced exon. RI: Retained intron. MXE: Mutually exclusive exons. **b.** PCA of exon usage matrices (Percent Spliced In, PSI) calculated by rMATS across WT and ZMYND11 deficient NSCs (n=4 replicates for WT and -/-; n=8 replicates using two clones for +/-). **c.** Triple overlap of DAS events prediction (HISAT2-rMATS, STAR-rMATS and AltAnalyze) to extract high-confidence DAS events in ZMYND11 -/- NSCs. **d.** Scheme outlining extracting tissue specific PSI for 590 high-confidence DAS events from *VastDB* database (https://vastdb.crg.eu/). **e.** Heatmap of tissue-specific PSI values (row-wise scaled by Z-score transformation) for high-confidence DAS events (y axis) in ZMYND11 -/- NSCs. 506 events with available tissue-specific PSI values are shown (1 event with the same values across tissues is omitted). Dividing line separates events that were either high in non-brain tissues (inclusion) or low in non-brain tissues (exclusion). Splicing pattern in ZMYND11 -/- NSCs was mapped to the heatmap on right side bar (red: inclusion events in KO; blue: exclusion events in KO). Numbers in red or blue show events whose splicing trends were matched between ZMYND11 -/- and non-brain tissues. **f.** Scatter plot depiction of PSI difference between WT and KO (x axis) & brain tissues and non-brain tissues (y axis). Red marks events with unpaired two-tailed *t* test *P*<0.05 between brain and non-brain tissue PSI, while black marks those with *P*>0.05. Closed circle marks events that showed matched trend between KO and non-brain tissues, while open circle marks those not matched. **g.** PSI fold changes relative to WT for all high-confidence DAS events in ZMYND11 -/- NSCs. Whiskers show min and max values. **h-i.** Representative high-confidence DAS events such as inclusion event *CTTN* exon 11 (**h**) and exclusion event *SYNGAP1* exon 14 (**i**) with Sashimi plots (upper), PSI changes (middle) and semi-quantitative PCR validation (lower). Numbers in Sashimi plots indicate average PSI values in WT or mutants. B: Brain. NB: Non-Brain. Statistics in **g**, **h**, **i**: one-way ANOVA followed by Tukey’s test for three groups, unpaired two-tailed *t* test for two groups. ns *P* > 0.05, \**P*<0.05; \*\**P*<0.01; \*\*\**P*<0.001; \*\*\*\**P*<0.0001. *P* values in **g**: *P*<0.0001. *P* values in **h**: +/- vs +/+: *P*=0.2034, -/- vs +/+: *P*<0.0001, -/- vs +/-: *P*=0.0003, NB vs B: *P*=0.0044. *P* values in **i**: +/- vs +/+: *P*=0.0035, -/- vs +/+: *P*<0.0001, -/- vs +/-: *P*=0.0014 NB vs B: *P*<0.0001.

To investigate the potential link between expression changes and DAS events, we examined tissue-specific percent spliced in (PSI) values from *VastDB*^22^ (**Fig. 4d**). We successfully mapped tissue-specific PSI values for 507 out of 590 DAS events identified in ZMYND11 -/- NSCs. Our analysis revealed a strong inverse relationship between PSI values in brain and non-brain tissues (**Fig. 4e**). Interestingly, for splicing events characterized by exon inclusion in non-brain tissues, the majority also showed exon inclusion in ZMYND11 -/- NSCs (205 out of 273 events). Conversely, for events typically exhibiting exon exclusion in non-brain tissues, a similar pattern was observed in ZMYND11 -/- NSCs (140 out of 233 events). Overall, approximately 70% of the DAS events in ZMYND11 -/- NSCs demonstrated a bias toward non-brain tissue splicing patterns. To further validate this observation, we correlated PSI values from ZMYND11 -/- cortical NSCs with non-brain tissue PSI values, finding a moderate correlation (R² = 0.23, **Fig. 4f**). In contrast, this correlation was nearly absent in ZMYND11 +/- NSCs (R² = 0.002, **Extended Data Fig. 7h**), suggesting that the loss of ZMYND11 disrupts brain-specific splicing, shifting it toward a non-brain splicing profile.

We further examined exon inclusion and exon exclusion DAS events in ZMYND11 -/- NSCs separately and found a dose-dependent PSI change (**Fig. 4g**). We validated high confidence exon inclusion and exclusion events using semiquantitative PCR (**Fig. 4h, i and Extended Data Fig. 8a, b**). We then compared our data with previously reported DAS events from ZMYND11 knockdown studies^5,6^, finding no significant overlap between these datasets (**Extended Data Fig. 7i**), which highlights the cell- and tissue-specific nature of alternative splicing regulation by ZMYND11.

To explore if ZMYND11 chromatin binding is associated with these alternative splicing changes, we identified direct binding to 68 out of 590 (∼11%) high confidence DAS events in ZMYND11 -/- NSCs (**Extended Data Fig. 7j**). This number increased to 141 events upon profiling FLAG-ZMYND11 binding (**Extended Data Fig. 7k**). Interestingly, while not all DAS events exhibited direct ZMYND11 binding, majority (∼70%) were located near the transcription start site (TSS +/- 1kb, **Extended Data Fig. 7l**), suggesting a potential role for ZMYND11 binding in regulating alternative splicing. Additionally, ZMYND11 binding was observed in both exon inclusion events (e.g., *APP*) and exon exclusion events (e.g., *L1CAM*, **Extended Data Fig. 7m**). This analysis suggests that ZMYND11 enables a switch in brain-specific alternative splicing.

### ZMYND11 deficiency causes a tissue-specific isoform switch that affects migration and proliferation of cortical NSCs

Mapping the differentially spliced isoforms to biological processes enriched for genes important for synapse organization and microtubule-based movement suggesting regulation of cytoskeletal machinery (**Extended Data Fig. 9a**). Specific exon usage in some of genes with DAS events have been previously linked to cancer cell migration^23–25^. To assess how these splicing changes might affect the migration of cortical NSCs, we designed shRNA constructs targeting either the brain or non-brain isoforms of *L1CAM*, *CTTN* and *MEAF6* (**Fig. 5a and Extended Data Fig. 9b**) as these genes exhibited high differential PSI values, ZMYND11 dose dependency, and displayed significant expression. For this experiment, we opted to use a stable NSC cell line (LTNSCs), which retain their progenitor state for over 100 passages while maintaining their neurogenic potential^26,27^. By using these isoform-specific constructs, we selectively targeted the brain or non-brain isoforms (**Extended Data Fig. 9c**) and performed scratch assays on a confluent monolayer of LTNSCs to determine the effects on cell migration. While we did not observe migration defects associated with knockdown of *CTTN* isoforms, targeting the brain isoform of *L1CAM* (exon 3 or exon 28) impaired LTNSC migration compared to control (**Fig. 5b**). Contrary to our predictions, the knockdown of the non-brain isoform of *MEAF6* not only blocked migration but also showed signs of low proliferation. To explore this further, we examined the cell cycle profiles of *MEAF6* brain versus non-brain isoforms in LTNSCs. BrdU labeling demonstrated that the knockdown of the non-brain isoform of *MEAF6* greatly decreased S phase (**Fig. 5c**). These results suggest that isoform switches impact both migration and cell proliferation in cortical NSCs.

**Fig. 5.**
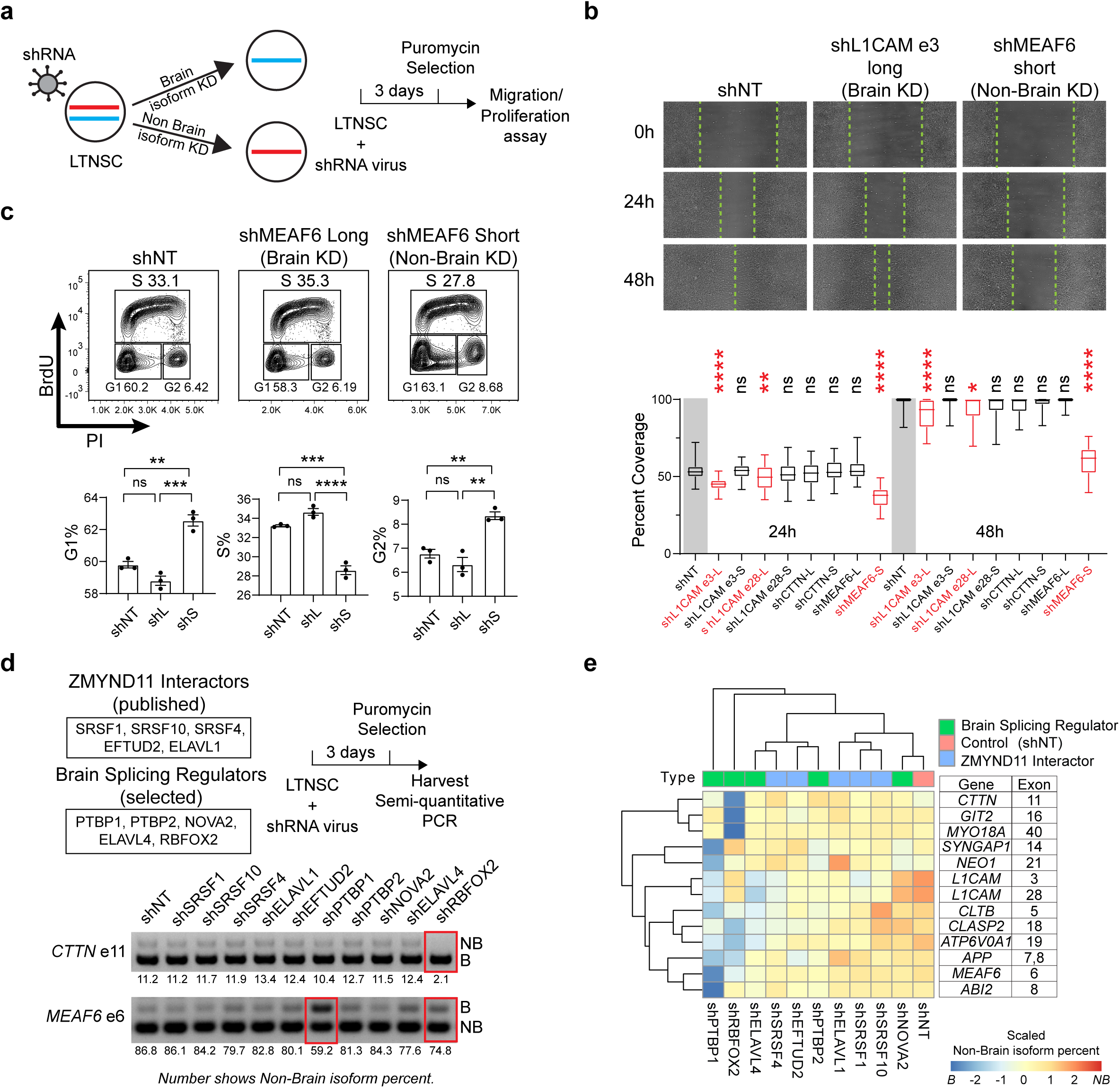
Non-brain-like isoforms alter migration and proliferation of cortical NSCs. **a.** Scheme outlining functional assays after specific isoform knockdown. Long-term neural stem cells (LTNSCs) were infected with short hairpin RNA lentiviruses to knockdown either brain isoform or non-brain isoform before puromycin selection. Then cells were used for either migration assay or proliferation assay. **b.** Migration assays (0h, 24h, 48h) on LTNSC with different isoform knockdown (upper) and quantification of scratch coverage compared with shNT (lower, n=48 different areas on 3 independent scratches for each knockdown experiment). Dashed lines mark migration edges in different time points. Red marks significantly changed shRNA. Whiskers show min and max values. NT: non targeting. **c.** BrdU-PI assays on LTNSCs with either MEAF6 long isoform (Brain isoform) or short isoform (Non-Brain isoform) knockdown (upper) with quantification (lower, mean±SEM, n=3 infections). **d.** Scheme outlining splicing factor screening experiment design (upper) and semi-quantitative PCR gels on *CTTN* exon 11 and *MEAF6* exon 6 (lower). LTNSCs were infected with shRNA lentiviruses that target either previously published ZMYND11 interactors or known brain splicing regulators before puromycin selection. The cells were then harvested for semi-quantitative PCR. Red rectangles mark significantly changed shRNA. Values shown below the gel are non-brain isoform percent. **e.** Heatmap for **d** using non-brain isoform percent values (row-wise scaled by Z-score transformation) on selected high-confidence DAS events that showed match between non-brain tissues and ZMYND11 KO. Note that *APP* has two DAS events that are combined as these two exons are close to each other. Statistics in **b**: one-way ANOVA followed by Dunnett’s test. Statistics in **c**: one-way ANOVA followed by Tukey’s test. ns *P* > 0.05, \**P*<0.05; \*\**P*<0.01; \*\*\**P*<0.001; \*\*\*\**P*<0.0001. *P* values in **b**: shL1CAM_e3-L vs shNT at 24h: *P*<0.0001, shL1CAM_e3-S vs shNT at 24h: *P*=0.9996, shL1CAM_e28-L vs shNT at 24h: *P*=0.0087, shL1CAM_e28-S vs shNT at 24h: *P*=0.3781, shCTTN-L vs shNT at 24h: *P*=0.2084, shCTTN-S vs shNT at 24h: *P*=0.9971, shMEAF6-L vs shNT at 24h: *P*=0.9786, shMEAF6-S vs shNT at 24h: *P*<0.0001, shL1CAM_e3-L vs shNT at 48h: *P*<0.0001, shL1CAM_e3-S vs shNT at 48h: *P*=0.9996, shL1CAM_e28-L vs shNT at 48h: *P*=0.0151, shL1CAM_e28-S vs shNT at 48h: *P*=0.4839, shCTTN-L vs shNT at 48h: *P*=0.6008, shCTTN-S vs shNT at 48h: *P*=0.9888, shMEAF6-L vs shNT at 48h: *P*=0.9575, shMEAF6-S vs shNT at 48h: *P*<0.0001. *P* values in **c**: shMEAF6-L vs shNT for G1%: *P*=0.1074, shMEAF6-S vs shNT for G1%: *P*=0.0012, shMEAF6-L vs shMEAF6-S for G1%: *P*=0.0002, shMEAF6-L vs shNT for S%: *P*=0.0543, shMEAF6-S vs shNT for S%: *P*=0.0002, shMEAF6-L vs shMEAF6-S for S%: *P*<0.0001, shMEAF6-L vs shNT for G2%: *P*=0.3799, shMEAF6-S vs shNT for G2%: *P*=0.0055, shMEAF6-L vs shMEAF6-S for G2%: *P*=0.0015.

While ZMYND11 expression is high in the brain, it is not exclusive. To further understand how ZMYND11 regulates brain-specific splicing, we tested the roles of several RNA-binding proteins (SRSF1, SRSF10, SRSF4, EFTUD2, ELAVL1) that have been reported as ZMYND11 interactors^5^ and selected brain-specific splicing regulators (PTBP1, PTBP2, NOVA2, ELAVL4, RBFOX2)^28–30^ using shRNA knockdown (**Extended Data Fig. 9d**). Semi-quantitative PCR analysis of 14 high-confidence DAS events revealed that only *PTBP1* and *RBFOX2* depletion resulted in a clear loss of non-brain isoforms (**Fig. 5d, e**). Interestingly, *RBFOX2* expression was slightly increased in *ZMYND11* mutants (**Extended Data Fig. 9e**). In contrast, knocking down other previously identified ZMYND11-interacting RNA-binding proteins did not significantly impact splicing, indicating that the splicing changes observed may involve multiple regulatory mechanisms. Finally, we explored whether BMP signaling might contribute to the observed splicing changes. We found that only high doses of BMP4 induced significant splicing changes, while low-dose BMP4 or BMP inhibitor treatments did not (**Extended Data Fig. 9f, g**). This suggests that despite ZMYND11 deficiency causing elevated expression of BMP signaling, it plays a limited role in regulating splicing in ZMYND11 knockout cortical NSCs.

### Tissue-specific isoform switches found in additional risk factor lines for ASD can be improved by ZMYND11

Our data thus far have implicated that ZMYND11 acts as both a regulator of latent gene activity and alternative splicing to promote faithful cortical development by acting as a repressive blanket over the epigenome. Due to this shared phenotype with other autism related risk factors (Class I ASD), we performed a similar set of experiments where we generated transcriptomic data derived from cortical NSCs (**Fig. 6a**). Expectedly, Class I ASD lines showed downregulation of genes linked to neurogenesis driving the separation with isogenic controls within the PCA plot (**Fig. 6b and Extended Data Fig. 10a**). Despite little connection with ZMYND11, Class I ASD lines appeared to upregulate the expression of developmental pathways including BMP and WNT compared to controls (**Fig. 6c**). Intriguingly, although lower in magnitude, we observed 20 significant high confidence DAS events that switched isoform tissue-specificity for which seven are shared with *ZMYND11* mutants (**Fig. 6d-f and Extended Data Fig. 10b**). Interestingly, this non-brain isoform trend was not observed in Class II ASD lines (**Fig. 6g and Extended Data Fig. 10c**). Since we found this unexpected similarity in the Class I ASD lines, we thought that ZMYND11, through its transcriptional repression function, could rectify the cortical NSC deficits by targeting the silencing of developmental pathways as well as promoting a brain-biased switch in isoform expression. We infected our dox inducible FLAG-ZMYND11 virus and differentiated individual Class I ASD lines as monolayer to cortical NSCs and assessed IPC generation (**Fig. 6h**). We found that while the percentage of IPC generation varied due to specific ASD mutations, there was indeed a positive correlation in the dox treated groups (**Fig. 6i, j**). These data suggest that both repression of latent developmental pathways and restoration of brain-biased isoforms aid in faithful corticogenesis and potentially implicate the activity of ZMYND11 or genes with similar function as critical for maintaining a stable cortical progenitor identity. To explore this idea, we overexpressed ZMYND11 in WT cortical NSCs for 72 hours to determine which gene regulatory networks are susceptible from being silenced (**Extended Data Fig. 10d**). Unexpectedly, transcriptome analysis of ZMYND11 overexpression in control lines revealed only ZMYND11 was significantly upregulated, suggesting that the homeostatic level of this chromatin reader was already at optimal levels, resulting in no significant impact on gene expression (**Extended Data Fig. 10e**).

**Fig. 6.**
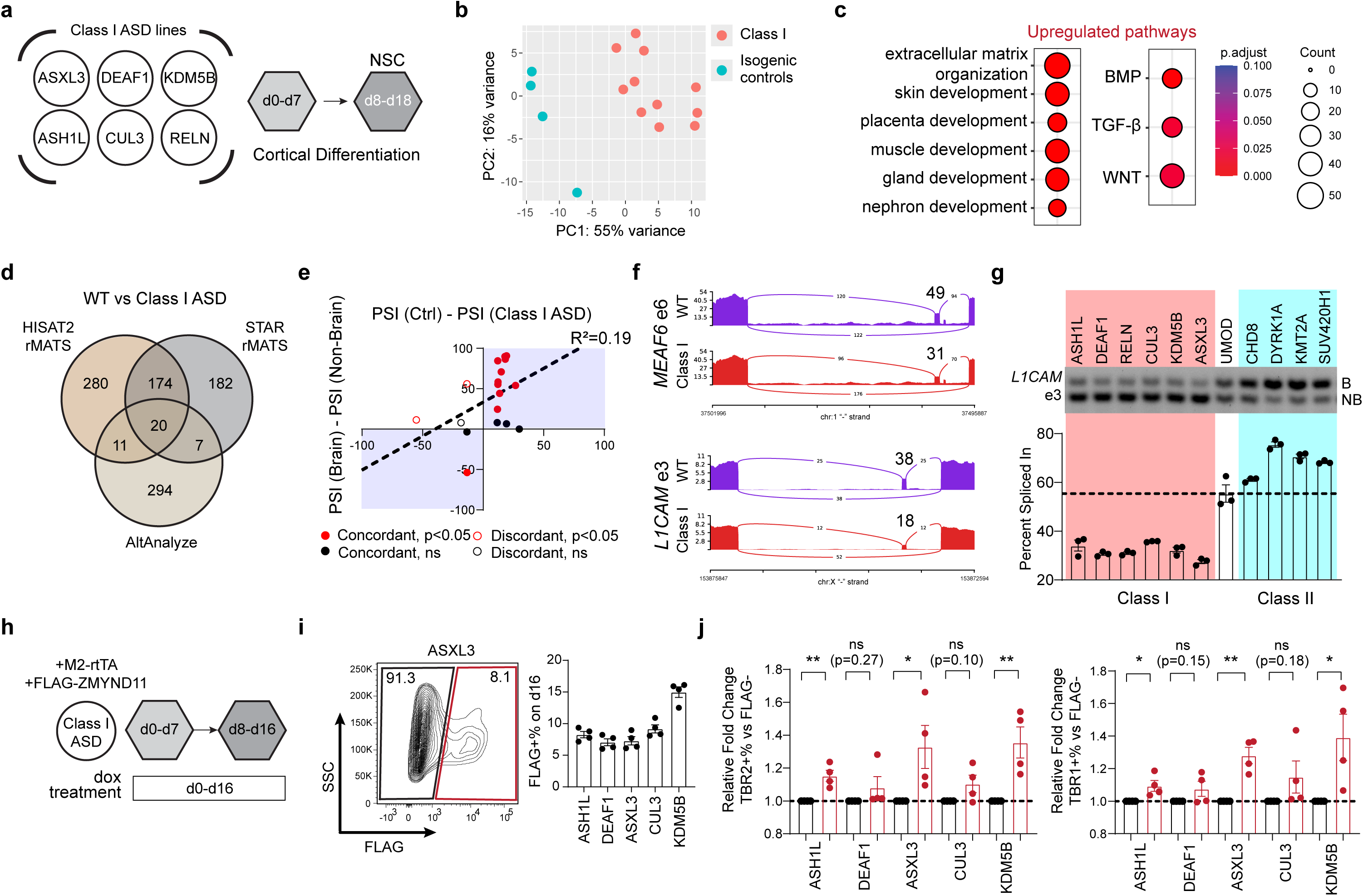
Several ASD risk factors share impaired IPC production and non-brain-like alternative isoform switches, which can be improved by ZMYND11. **a.** Scheme outlining Class I mutant lines and cortical differentiation. **b.** Principal component analysis of RNA-seq on Class I mutant and isogenic control cortical NSCs (n=11 for Class I mutants, n=4 for isogenic controls). **c.** Gene ontology analysis on upregulated genes in Class I NSCs using biological processes. *P*.adjust values were calculated using Benjamini-Hochberg method. **d.** Triple overlap of DAS events prediction (HISAT2-rMATS, STAR-rMATS and AltAnalyze) to extract high-confidence DAS events in Class I mutant NSCs. **e.** Scatter plot depiction of PSI difference between isogenic control and Class I mutants (x axis) & brain tissues and non-brain tissues (y axis). **f.** Sashimi plots for *MEAF6* exon 6 (upper) and *L1CAM* exon 3 (lower) from high-confidence DAS events in Class I mutant NSCs. Numbers in Sashimi plots indicate average PSI values. **g.** Semiquantitative PCR of *L1CAM* exon 3 on both Class I mutant and Class II mutant NSCs and quantification (mean±SEM, n=3 differentiations). **h.** Scheme outlining ZMYND11 overexpression strategy on Class I mutant differentiations. **i.** Intracellular flow cytometry analysis for FLAG on dox-treated (200ng/ml) Class I mutant monolayer culture at day 16 (left) with quantification (right, mean±SEM, n=4 differentiations). **j.** Quantification of intracellular flow cytometry analysis for TBR2 (left) and TBR1 (right) on FLAG+ populations at d16 relative to FLAG-population (mean±SEM, n=4 differentiations). Statistics in **j**: unpaired two-tailed *t* test. ns *P* > 0.05, \**P*<0.05; \*\**P*<0.01. *P* values in j: TBR2+% for *ASH1L*, *P*=0.0042; for *DEAF1*, *P*=0.2700; for *ASXL3*, *P*=0.0456; for *CUL3*, *P=*0.0998; for *KDM5B*, *P=*0.0094. TBR1+% for *ASH1L*, *P*=0.0303; for *DEAF1*, *P*=0.1519; for *ASXL3*, *P*=0.0013; for *CUL3*, *P=*0.1848; for *KDM5B*, *P=*0.0338.

From these data, we conclude that a subthreshold of ZMYND11 levels exhibits defective generation of IPCs from cortical NSCs, a critical cell type for mammalian neurogenesis. Upregulation of latent developmental related pathways and a switch from brain to non-brain alternative splicing impair the efficiency in generating these cells from cortical NSCs. Intriguingly, elevated levels of BMP and WNT signaling as well as the non-brain isoform switch is a shared phenotype found with mutants in other high-confidence ASD-risk genes whose corticogenesis can be improved by ZMYND11 overexpression. These findings highlight the complexity of chromatin-associated pathogenic variants where dysregulation of gene expression is not the sole cause of NDD/ASD.

## Discussion

Our study uncovers a pivotal role for ZMYND11 as a dual regulator of chromatin dynamics and splicing fidelity in cortical neural stem cells (NSCs), providing a novel lens through which to understand the molecular underpinnings of ASD and related NDD. We were able to correct isoform-specific splicing defects and restore neurogenesis to levels reflective of controls by modulating splicing factor expression or overactivation of ZMYND11. Leveraging this knowledge, we envision precision medicine strategies that may promote splicing modulation as a potential therapeutic route^31^.

An intriguing result was the abnormal activation of BMP and WNT signaling pathways in ZMYND11 deficient cortical NSCs. Prior studies have shown that these pathways play essential roles in regulating corticogenesis^32–35^, and abnormal WNT signaling has been proposed as a major dysregulated pathway conferring risk for NDD/ASD pathogenesis^36,37^. In agreement with this, ZMYND11 deficient NSCs displayed elevated levels of BMP & WNT signaling associated with increased expression of genes related to non-neural development. We found that ZMYND11 directly binds these genes to repress their activation, although unexpectedly we noticed that ZMYND11 binding could not be fully explained by H3K36me3 enrichment. One candidate factor could be H3.3 S31ph which has been shown to eject ZMYND11^17^. Notably, while BMP inhibition rescued IPC production, WNT inhibition did not, likely because WNT signaling plays a more significant role in regulating anterior-posterior identity which was not explored in our monolayer culture system^38^. Future studies could explore the contribution of non-canonical WNT signaling, particularly WNT5A, which was also highly elevated in ZMYND11 deficient lines^39^.

One of the most striking findings from our study identified splicing alterations as a critical mechanism in ZMYND11 deficient cortical NSCs. Alterations in splicing, specifically, in ‘microexons’ have been previously reported as enriched in patients with ASD^20,28,40^. ZMYND11 caused splicing changes that biased toward non-brain tissue identity reflected a broader gene expression shift toward activation of non-neural-like development. Importantly, this switch in alternative splicing isoforms are not merely byproducts but actively contribute to cellular deficits, affecting processes like migration and proliferation in NSCs. Surprisingly, we found that ZMYND11 did not largely regulate splicing, as few DAS events overlapped with chromatin binding. This suggests that the dysregulation of splicing is likely indirect, potentially mediated by splicing genes such as RBFOX2^41,42^, since its expression was increased in *ZMYND11* mutants. Overall, our findings point to a multi-faceted regulation of splicing by ZMYND11, with both direct and indirect mechanisms at play.

Although multiple studies have identified different convergent mechanisms in NDD/ASD pathogenesis^43–47^, most studies are limited in phenotypic convergence or face problems such as low-depth sequencing. Moreover, as the idea of non-neural fate shift has also been previously reported in ASD cortex where there was an enrichment of non-neural microexon^20^ and in an additional NDD/ASD risk gene, *MYT1L*, whose mutants elevates gene expression of non-neural fate and impair neurogenesis^48,49^, it remains to be determined whether the same concept applies to a broader group of NDD/ASD mutations. To this end, we differentiated cortical NSCs from 6 separate lines harboring risk factors for *ASH1L, DEAF1, RELN, CUL3, KDM5B, ASXL3* along with multiple isogenic controls and revealed similar transcriptional changes as those found in ZMYND11 that included the impairment of IPC generation and upregulation of latent developmental pathways. Although splicing alterations in these lines were less pronounced, they exhibited a similar trend toward non-brain tissue bias potentially due to the greater sequencing depth we used. Interestingly, this non-brain splicing trend did not apply to all NDD/ASD risk genes, as Class II mutants (ones that enhance neurogenesis) showed an opposite splicing trend. This suggests that NDD/ASD encompasses not just a spectrum of phenotypes but a range of underlying molecular mechanisms that correlate closely with the biological outcomes. Nevertheless, strategies to modulate RNA splicing that have already been successful in spinal muscular dystrophy treatment^50^ serve as a novel therapeutic avenue for Class I mutants.

The strive for discovering common molecular mechanisms in polygenic disorders is to develop broad therapeutic strategies, reducing the need for personalized interventions. As a proof of principle, we demonstrated that overexpressing ZMYND11 in Class I mutant lines partially rescued cortical NSCs deficits by suppressing non-neural gene expression and restoring brain-biased splicing. Importantly, ZMYND11 overexpression in WT NSCs did not induce adverse effects, suggesting that ZMYND11 levels are not dose sensitive. This feature, coupled with ZMYND11’s relatively small size compared to other Class I genes (e.g., ASH1L, KDM5B, ASXL3), makes it a promising candidate for gene therapy approaches for which multiple studies have tried overexpression as a gene therapy^51,52^. Future work will examine how overexpressing ZMYND11 partially restores epigenetic landscape in Class 1 mutants by potentially compensating for other epigenetic gene dysregulation.

## Methods

### Maintenance and propagation of human embryonic stem cells (hESCs)

Human embryonic stem cells (hESCs) were maintained in Essential 8 or Essential 8 flex (E8) medium on Vitronectin-coated dishes following established protocols. Briefly, hESCs (H1, XY; H9, XX; MEL1, XY) were cultured at 37°C in a 5% CO_2_ environment and passaged as clumps using an EDTA dissociation solution. Cells were used for differentiation between passage 50-70. All hESC lines are karyotypically normal and tested for mycoplasma every 2-3 months.

### Human embryonic kidney 293T (HEK293T) cell culture and lentivirus production

HEK293T cells were cultured at 37°C in a 5% CO_2_ environment using DMEM medium supplemented with 10% fetal bovine serum (FBS), 1X Antibiotic-Antimycotic and 1X GlutaMAX. The medium was changed every 2 days, and cells were passaged using Trypsin-0.05% EDTA.

Lentiviruses were produced as previously described. The day before transfection, 5 million HEK293T cells were plated in a 10 cm dish coated with Poly-L-lysine & 0.1% gelatin. The lentivirus, packaging (psPAX2) and envelope (pMD2.G) vectors were transfected using X-tremeGene 9 in a 1:2:1 molar ratio, respectively. For the M2-rtTA and FLAG-ZMYND11 vectors, medium was changed into E8 18 hours post transfection. For the pGreenPuro shRNA vectors, medium was changed into N2 medium supplemented with B27 (1:1000, without Vitamin A; N2-B27), 20 ng/ml EGF and 20ng/ml FGF2 18 hours post transfection. Viruses were harvested at 48 and 72 hours post transfection, concentrated using AMICON Ultra-15 Centrifugal Filter Units, and snapped frozen in liquid nitrogen.

### Three-dimensional guided brain organoid differentiation

Guided cortical organoid differentiation involved seeding 5000 hESCs into one well of ultra-low attachment 96 well plate in E8 supplemented with 10 μM ROCKi on day −1. EBs were treated with 100 nM LDN193189 (BMPi), 10 μM SB431542 (TGFβi) and 5 μM XAV939 (WNTi) in Essential 6 (E6) medium from day 0 to 6. The medium was then replaced with N2 medium supplemented with 20 ng/ml EGF, 20 ng/ml FGF2, and 1:1000 of B27 supplement without RA from day 7 to day 19. Cortical Spheroids were further matured in 20 μg/ml BDNF, 20 μg/ml GDNF and 200 μM Ascorbic acid in Neurobasal medium supplemented with B27 (1:50, without Vitamin A; NB-B27) and 1X GlutaMAX for up to 30 days.

### Monolayer prefrontal (PFC) cortical differentiation and excitatory neuron induction

Dishes were coated with Matrigel (diluted 1:100 in DMEM/F12) and stored overnight at 4°C. The next day, hESCs were dissociated to single cells using Accutase, resuspended with E8 and plated on the Matrigel coated dish at a density of 300,000 cells /cm^2^ in the presence of 10 μM ROCKi on day −1. From day 0 to 2, the cells were cultured in E6 medium supplemented with 100 nM LDN193189, 10 μM SB431542 and 5 μM XAV939 (i.e., LSBX) to promote anterior telencephalic patterning. From day 3 to 7, the cells were maintained in E6 with LSB with XAV939 removed. From day 8 to 18, the cells were treated with 50 ng/ml FGF8 in N2-B27 to promote a PFC-like cortical regionalization.

To induce cortical NSCs to neurons, 10 μM DAPT (NOTCHi) was added to day 18 of differentiation in NB-B27 for 5 days. The neurons were then replated onto dishes coated with Poly-ornithine (PO), Laminin (Lam) and Fibronectin (FN) and maintained in NB-B27 supplemented with 20 μg/ml BDNF, 20 μg/ml GDNF and 200 μM Ascorbic acid. To eliminate proliferative cells from the neuron culture, 1 μM Ara-C was added for 48 hours. Fifty percent of neuron medium was changed every 3 days.

### Maintenance of long-term neural stem cells (LTNSCs)

Derivation of LTNSCs was as previously described^26,27^. LTNSCs were maintained on PO-Lam-FN coated dishes with 20 ng/ml EGF and 20 ng/ml FGF2 in N2-B27. LTNSCs were maintained at high density, used between passage 15-25, and passaged every week using 0.05% Trypsin.

### Spontaneous differentiation of hESCs

5000 hESCs were seeded into one well of ultra-low attachment 96 well plate in E8 supplemented with 10 μM ROCKi on day −1. From day 0 to 10, EBs were treated 20%KSR, 3%FBS, 1X GlutaMAX and 1X MEM-NEAA in DMEM/F12 medium before harvest at day 10.

### Plasmid constructs and Molecular cloning

The lentiviral vector used to overexpress human ZMYND11 in our study, pLV[Exp]-Puro-TRE>3xFLAG-hZMYND11[NM_001370102.2], was constructed by VectorBuilder. The vector ID is VB211020-1233tqt, which can be used to retrieve detailed information about the vector on vectorbuilder.com.

Short hairpin RNA (shRNA) sequences were manually designed and cloned into the restriction sites EcoRI & BamHI in the pGreenPuro vector (System Biosciences, SI505A-1). shRNA duplexes were annealed and inserted using traditional ligation cloning. All shRNA sequences were listed in **Supplementary Table 4**.

### Generation of *ZMYND11* mutant hESCs

CRISPR/Cas9 was used to introduce frameshift mutations into exon 3 of ZMYND11 in H1 hESCs. Guide RNA (gRNA, IDT): 5’-AGCTAAGCTCAGCTGACGGG was chosen using predictive scores from website tool (http://crispor.tefor.net/). The gRNA-Cas9 complex was transfected using Lipofectamine Stem Transfection Reagent (Invitrogen) according to manufacturer’s protocol. Individual clones were isolated by replating transfected cells at low density. Indels were screened using Sanger sequencing, and heterozygous and compound heterozygous clones were inferred bioinformatically using DECODR (https://decodr.org). To identify mutations on different alleles and to exclude mixed clones, TOPO cloning of the targeted region was applied. Selected heterozygous clones (e.g., Het2, Het4) were clonally expanded a second time to ensure there was no mixing wildtype and homozygous clones.

### Generation of doxycycline inducible ZMYND11 overexpression lines

In experiments where we overexpressed the wildtype ZMYND11 construct, PX458-AAVS1 (Addgene #113194) and AAVS1-Neo-M2rtTA (Addgene #60843) were nucleofected into 3 million cells by LONZA 4D-Nucleofector using program CB-150. After 3 days of recovery with CloneR, the cells were selected using 200 μg/ml Geneticin and replated at low density for clonal expansion. Single colonies were isolated and verified for biallelic targeting by genomic PCR.

To overexpress ZMYND11 in ZMYND11 knockout or wildtype hESCs, validated and karyotypically normal M2-rtTA hESC clones were infected with 3X-FLAG-ZMYND11 lentivirus, followed by the addition of 2 μg/ml puromycin after 72 hours to select for resistant clones. For overexpression of ZMYND11 in Class 1 gene mutant lines, FUW-M2rtTA (Addgene #20342) and 3X-FLAG-ZMYND11 lentiviruses were both transduced, with puromycin selection applied to isolate resistant clones.

### Intracellular FACS analysis

Cells were dissociated using Accutase and washed with DPBS. Approximately 5 x 10^5^ cells were treated with Zombie Violet Live/Dead Dye (1:1000) in DPBS for 30 minutes. After washing with DPBS, the cells were stained with surface antibodies for 30 minutes (if needed). Next, the cells were fixed using 1X Fix/Perm buffer from the BD Pharmingen Transcription Factor Buffer Set and incubated at 4°C for 1 hour. Cells were then washed twice with 1X Perm/Wash buffer and incubated with primary antibodies for 1 hour. After 2 more washes, secondary antibodies were added, and cells were incubated at 4°C for 1 hour in the dark. Finally, the cells were washed 2 times and prepared for analysis.

### Cell cycle exit analysis

Cell cycle analysis was performed according to manufacturer’s protocol. Briefly, cells were treated with 10μM BrdU for 30 minutes and harvested using Accutase. The cells were then fixed with 70% ethanol on ice for 30 minutes, followed by treatment with 2N HCl/Triton X-100 at room temperature for 30 minutes, and neutralized with 0.1M borate buffer. After washing, the cells were stained with FITC anti-BrdU or AF647 anti-BrdU for 30 minutes at room temperature. Cells were washed again and resuspended in 5 μg/ml propidium iodide and prepared for analysis.

### Histology and Immunohistochemistry

For 3D culture, organoids were fixed in 4% PFA overnight at 4°C, followed by three washes with DPBS the next day. The organoids were then dehydrated in 30% sucrose solution overnight at 4°C and sectioned at a 20-micron thickness. Sections were mounted onto glass slides, blocked with 5% Normal Donkey Serum (NDS) for 30 minutes and incubated with primary antibodies overnight at room temperature. Sections were then washed 3 times with 0.2% Tween-20 and incubated with secondary antibodies for 2 hours at room temperature. The sections were then washed again, stained with DAPI, embedded in Fluoromount, covered with coverslips, and stored at 4°C until imaging.

For 2D culture, cells were replated onto glass coverslip 24 hours before fixation. The cells were fixed in 4% PFA for 10 minutes at 4°C, permeabilized with 0.5% Triton X-100 for 5 minutes and blocked with 5% NDS in 0.2% Tween-20 for 1 hour. The cells were then incubated with primary antibodies overnight at 4°C. The cells were then washed 3 times with 0.2% Tween-20 and incubated with secondary antibodies for 30 minutes at room temperature, washed and stained with DAPI. The cells were then embedded in Fluoromount and stored at 4°C until imaging.

### Western blot

Cells were dissociated and counted. An equal number of cells were harvested per experiment, lysed in 1X LDS Sample Buffer supplemented with 1X Sample Reducing Agent then denatured at 95°C for 5 minutes. Samples were then loaded and run on a 4-12% pre-cast Bis-Tris gel and transferred onto Nitrocellulose membrane for 2 hours. Membranes were blocked in 5% milk and primary antibodies were incubated overnight at 4°C. The next day membranes were washed with 1X TBS buffer with 0.1% Tween-20 and incubated with HRP-conjugated secondary antibodies for 1 hour. Blots were visualized using SuperSignal Chemiluminescent Substrate.

### RNA extraction and qPCR

RNA was extracted using TRIzol followed by chloroform extraction. Purification of RNA was performed using RNeasy Kits and eluted in nuclease free water. cDNA synthesis was performed with 1 μg of RNA using High-Capacity cDNA Reverse Transcription Kit. qPCR was performed with SYBR Green using QuantStudio 3 Real-Time PCR system. For semi-qPCR, isoform specific primer pairs were designed and used to amplify both the inclusion and exclusion products. Products were resolved on high percentage (3%) agarose gel electrophoresis. Calculation of percent spliced in was done by measuring the intensity of inclusion isoform band divided by the sum of intensity of inclusion and exclusion isoform bands using ImageJ. Calculation of non-brain isoform percent was done similarly by measuring the intensity of non-brain isoform band divided by the sum intensity. All primer sequences were listed in Supplementary Table 4.

### Scratch/Migration assay

LTNSCs were seeded onto 6-well plates at density of 3 x 10^5^ cells/cm^2^ and infected with shRNA viruses the next day before puromycin selection. After the cells became 100% confluent, a scratch was performed on the monolayer using a sterile 200 μl micropipette tip. The cells were carefully washed with the medium to remove non-adherent cells, and cultured for an additional 2 days monitoring cell migration every 24 hours. The rate of migration was measured by quantifying the distance that the cells moved from the edge of the scratch to the center as previously described^53^.

### RNA-seq and gene expression analysis

Total RNA for hESCs was poly-A enriched using oligodT Dynabeads (Invitrogen) and strand-specific RNA-seq libraries were constructed using TruSeq Stranded mRNA library kit, following the manufacturers’ instructions. Libraries were submitted to Roy J. Carver Biotechnology Center at the University of Illinois at Urbana-Champaign (UIUC) for paired-end sequencing (150bp). Total RNA for cortical neural stem cells was submitted to Novogene for paired-end sequencing (poly-A enriched, 150bp). Raw FASTQ files were quality checked using FastQC. Adapter sequences were trimmed using fastp with parameters “-q 20 -n 7”. Trimmed reads were then aligned to ENSEMBL GRCh38 using HISAT2 with default parameters. A raw count file was generated using featureCounts in Subread with default parameters and genes with no raw counts in >= 75% of samples were filtered out. Downstream analysis was done using R packages. Normalized counts per million (cpm) values were generated using “cpm” in “edgeR”. Principal component analysis was performed using “plotPCA” in “DESeq2”. Differential gene expression analysis was performed using “DESeq2” with *P*.adjust < 0.05 and fold change > 1.5. Gene ontology analysis was performed using “clusterProfiler” with *P*.adjust calculated using Benjamini-Hochberg method. Plots were generated using “ggplot2”.

### Alternative splicing analysis using rMATS

Input alignment files for rMATS splicing analysis were generated either using HISAT2 with trimmed reads or STAR with raw reads with default parameters. Results from rMATS were further filtered by removing low coverage events (avg_read >5) using “maser” in R. Significant splicing events were identified with FDR < 0.05 and percent spliced-in change (ΔPSI) > 0.1. Sashimi plots were generated using rmats2sashimiplot. For exon usage principal component analysis, PSI matrix was generated using rMATS with parameters “-- statoff”. PSI matrix was then imputed by KNN method for missing values using “impute” in R. PCA was conducted using “prcomp” function in R.

For RNA-seq datasets reanalysis in Guo et al., 2014 and Wen et al., 2014, we only used HISAT2-rMATS for the alternative splicing analysis.

### Alternative splicing analysis using AltAnalyze

For AltAnalyze we only input BAM files generated by STAR as indicated by user manual. Significant splicing events were identified with raw *P* value < 0.05 and ΔPSI > 0.1. Events from AltAnalyze were more than those from rMATS as AltAnalyze summarized events at each individual exon-exon or exon-intron junction while rMATS summarized events of individual exon. Therefore, a significant changed spliced exon or intron might have more than one event reported in AltAnalyze.

Overlapping significant spliced exon events by STAR-AltAnalyze with those from HISAT2-rMATS or STAR-rMATS was performed using genomic coordinates of the target exon. To expand the potential overlap, events that have matched upstream & downstream exon coordinates and either matched 5’ target exon coordinate or 3’ target exon coordinate were also included. Plus, events predicted from different methods would have to be in the same direction (both predicted to be spliced-in or spliced-out) to be considered as overlapped. These overlapped events were subsequently called high confidence differential alternative splicing (DAS) events for further analysis.

Tissue-specific PSI values for high confidence DAS events were extracted from *VastDB* database (https://vastdb.crg.eu) according to target exon coordinates in ENSEMBL GRCh38 and categorized into either brain tissue or non-brain tissue for comparison. Heatmap was generated using “pheatmap” in R after missing values were imputed by KNN method using “impute” in R. Correlation plots were generated using Prism.

### CUT&RUN, library preparation and analysis

CUT&RUN method was as previously described^54^. Briefly, 5 x 10^5^ cells were harvested and bound to Concanavalin A beads. These cell-bound beads were incubated with antibodies overnight at 4°C and treated with pAG-MNase for 1 hour at 4°C. Calcium was added to start pAG-MNase digestion for 30 minutes and CUT&RUN fragments were released and purified by phenol-chloroform extraction. DNA concentration was measured by Quantus fluorometer. For library preparation, we followed NEBNext Ultra II DNA Library Prep Kit for Illumina and performed size selection after PCR enrichment of adaptor-ligated library using MagBind Magnetic Beads. Libraries were sent to Novogene for sequencing aiming for 10-20 million reads.

CUT&RUN sequencing reads were quality checked using FastQC, and adapters were trimmed using fastp with parameters “-q 20 -n 7”. Reads were aligned to ENSEMBL GRCh38 using bowtie2 with parameters “--very-sensitive --no-unal --no-mixed --no-discordant -I 10 -X 700”. Low quality (MAPQ<10) reads were discarded using Samtools view with parameters “-q 10”. Duplicated reads were discarded using Picard MarkDuplicates with parameters “REMOVE_DUPLICATES=true”. Only uniquely mapped reads were used for subsequent analysis after these filters. Bigwig files were generated using bamCompare in deepTools with parameters “--binSize 20 --normalizeUsing RPKM”. For samples with replicates BAM files were merged before generating bigwig files. Visualization was performed using computeMatrix with parameters “scale-regions --beforeRegionStartLength 5000 --regionBodyLength 5000 --afterRegionStartLength 5000 --skipZeros -- missingDataAsZero” and plotHeatmap in deepTools. Spearman correlation was performed using multiBigwigSummary and plotCorrelation with default parameters in deepTools. Peak calling was performed using getDifferentialPeaksReplicates.pl in HOMER to call ZMYND11-bound regions versus ZMYND11 KO sample and FLAG-ZMYND11-bound regions versus no doxycycline sample with poisson *P* value < 0.05 and fold change >1. For ZMYND11 binding calling in FLAG-ZMYND11 overexpression experiment, we used rabbit IgG as the control and peaks were called using findPeaks.pl. H3K36me3 enriched peaks were called also against rabbit IgG control. Peaks overlapping with ENCODE blacklist regions were discarded. The blacklist file was downloaded from GitHub (https://github.com/Boyle-Lab/Blacklist/blob/master/lists/hg38-blacklist.v2.bed.gz). Overlapping of peaks was performed using bedtools intersect with default parameters. Annotation of peaks was performed using UROPA with parameters “feature: transcript, distance: 10000, attribute.value: protein_coding”. Clustering of ZMYND11-bound genes was done using plotHeatmap with parameters “--kmeans 3” in deepTools. Gene ontology analysis was similarly performed using “clusterProfiler” in R.

Re-analysis of previous ZMYND11 binding datasets were similarly performed using the same pipeline (mouse datasets mapped onto ENSMBL GRCm39, human datasets mapped onto ENSEMBL GRCh38) except during alignment without “-I 10 -X 700” parameters in bowtie2.

Overlapping of peaks with DAS events was similarly performed using bedtools intersect with default parameters. Distance of DAS events to TSS was calculated and plotted using “ChIPseeker” in R.

### Quantification, statistical analysis and reproducibility

All data represent mean ± SEM. Statistical significance was determined by parametric tests including unpaired two-tailed t test for two groups and analysis of variance (ANOVA) for more than two groups with post hoc tests as indicated. A *P* value <0.05 was considered statistically significant. *P.*adjust values in gene ontology analysis were calculated using Benjamini-Hochberg method. Prism 8 software was used for statistical analysis. For all cell culture experiments, n represents total number of independent differentiations. Each experiment was repeated at least 3 times.

## Data and Code Availability

The RNA and CUT&RUN sequencing data generated in this paper is uploaded to GEO with accession number GEO: GSE279072 and GSE279068, respectively. This dataset also includes raw count tables of RNA-seq and bigWig files of CUT&RUN-seq. This paper does not report original code.

## Acknowledgements

We thank Drs. Rupa Sridharan (University of Wisconsin, Madison), Kenneth Campbell (CCHMC) and members of the Tchieu lab for their critical reading, helpful critiques, and advice on this study. Additionally, we would like to thank Dr. Qing (Richard) Lu (CCHMC) for providing the pGreenPuro vector.

## Author contributions

Conceptualization: XC, JT

Methodology: XC, WL, SM, CH, GYC, LS, MI, AS, CC, JT

Investigation: XC, CC, JT

Visualization: XC, JT

Funding acquisition: JT

Project administration: JT

Supervision: CC, JT

Writing – original draft: XC, JT

Writing – review & editing: XC, WL, SM, CH, GYC, LS, MI, AS, CC, JT

## Data and materials availability

All data, code, and materials used in our analysis is available to any researcher for purposes of reproducing or extending the analysis. Cell lines can be provided through materials transfer agreements (MTAs). The RNA and CUT&RUN sequencing data generated in this paper is uploaded to GEO with accession numbers: GSE279072 and GSE279068, respectively. This dataset also includes raw count tables of RNA-seq and bigWig files of CUT&RUN-seq. This paper does not report original code.

## Supplementary Materials

Extended Data Figures 1 to 10.

Supplementary Tables 1 to 5.

---

**Extended Data Fig. 1.**
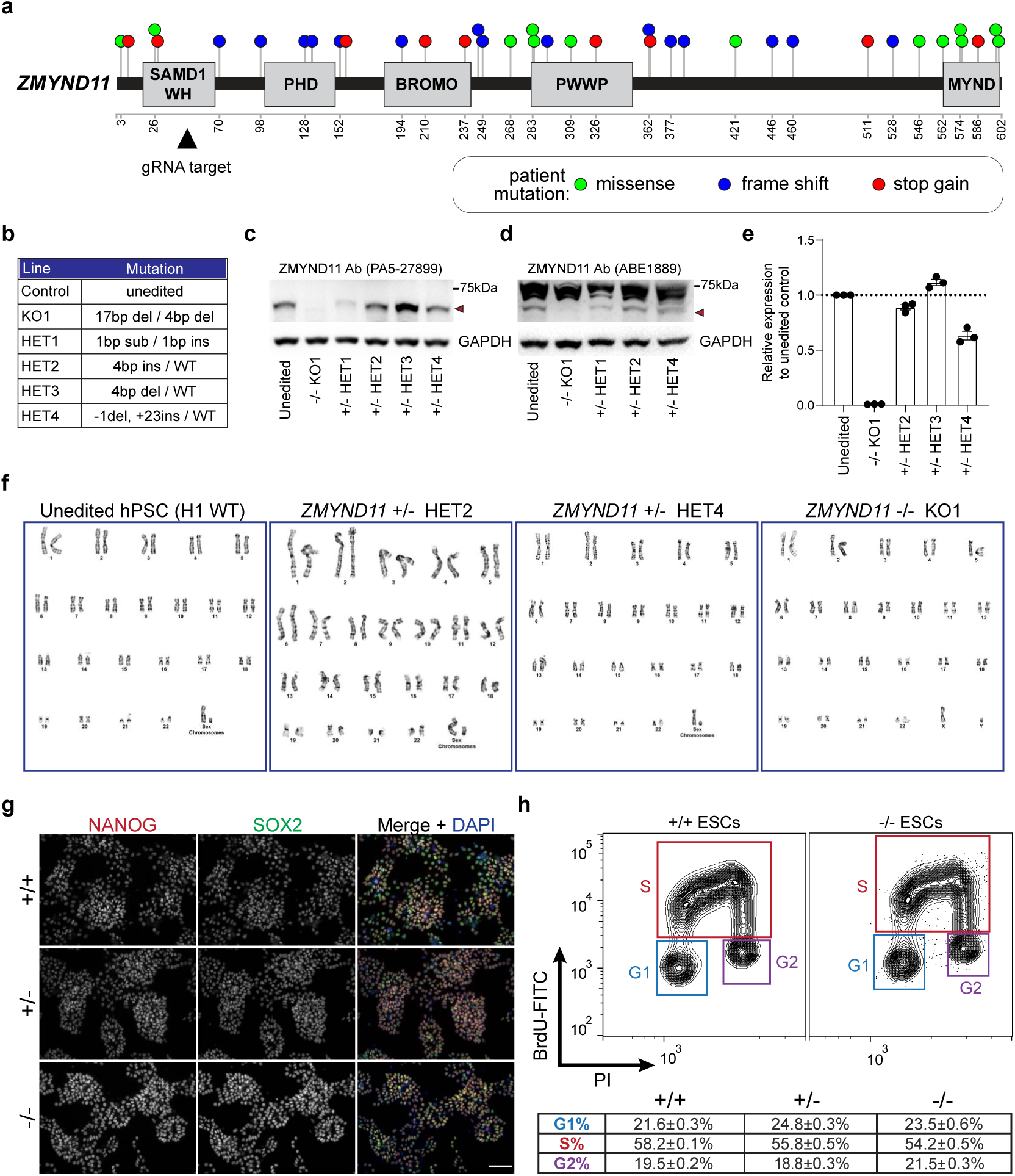
Engineering and characterization of *ZMYND11* mutant hESCs. **a**. Patient *ZMYND11* mutations distribution from SFARI database (https://gene.sfari.org/, data extracted in May 2024). 27/51 mutations are either stop-gained or frameshift mutations, 18/51 mutations are missense mutations, and 6/51 mutations are splice site mutations. gRNA was designed to target *ZMYND11* exon 3 (arrowhead). **b**. Genotypes of CRISPR-edited *ZMYND11* mutant clones in H1 hESC background. **c-d**. Western blot using two ZMYND11 antibodies (PA5-27899, ABE1889) on ZMYND11 deficient hESCs. **e**. Quantification of ZMYND11 western blot to confirm ZMYND11 knockout (mean±SEM, n=3). Het2 and Het4 were used for subsequent studies. **f**. Karyotype of unedited (WT) and ZMYND11 deficient clones (+/-, -/-). **g**. Immunofluorescence for pluripotency markers NANOG and SOX2 on WT and ZMYND11 deficient hESCs. Scale bars, 50μm. **h**. BrdU-PI cell cycle profiling on WT and ZMYND11 deficient hESCs (upper) and quantification (lower, mean±SEM, n=3 for WT and -/-, n=6 using two clones for +/-).

**Extended Data Fig. 2.**
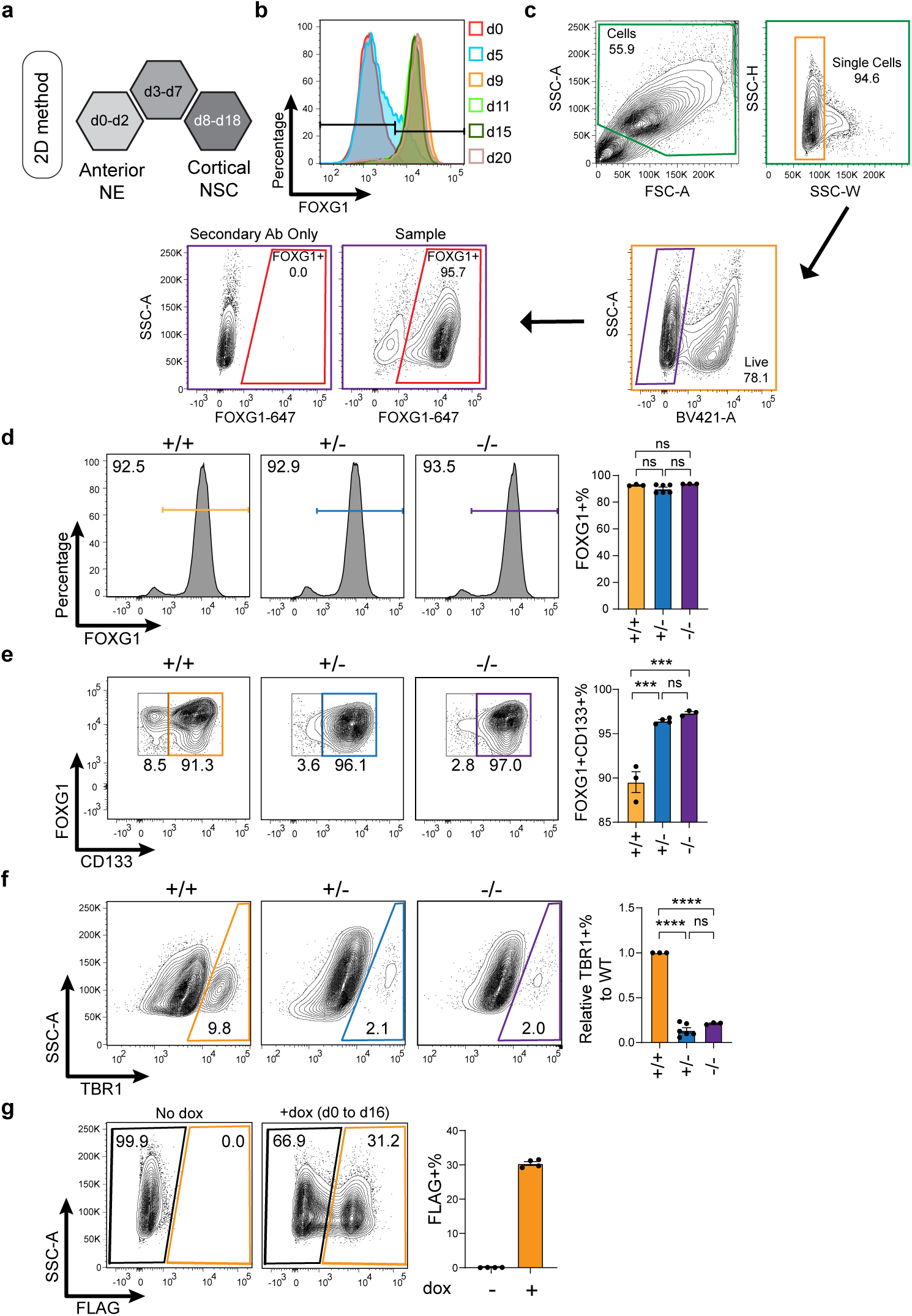
Quantitative analysis using intracellular flow cytometry on WT and ZMYND11 deficient monolayer culture. Related to. Figure 1**. a**. Schema outlining two-dimensional monolayer differentiation protocol. NE: neuroectoderm. **b**. Intracellular flow cytometry analysis for FOXG1 throughout early monolayer differentiation. Y axis shows percentage (“normalized to mode” option in FlowJo). **c**. Intracellular flow cytometry analysis gating strategy. Representative gating strategy for all panels: size selection, single cell selection, live cell selection, gates set on secondary antibodies stained only negative control. **d-f**. Intracellular flow cytometry analysis for FOXG1 (**d**), CD133 (**e**) and TBR1 (**f**) on WT and ZMYND11 deficient monolayer culture at day 16 (left) with quantification (right, mean±SEM; for **d**, n=3 differentiations for WT and -/-, n=6 differentiations using two clones for +/-; for **e**, n=3 differentiations for WT and -/-, n=4 differentiations using two clones for +/-; for **f**, n=3 differentiations for WT and -/-, n=6 differentiations using two clones for +/-). Y axis in **e** shows percentage (“normalized to mode” option in FlowJo). **g**. Intracellular flow cytometry analysis for FLAG on no doxycycline and d0-d16 dox-treated monolayer culture at day 16 (left) with quantification (right, mean±SEM, n=4 differentiations). Statistics in **d**, **e**, **f**: one-way ANOVA followed by Tukey’s test. ns *P* > 0.05, \**P*<0.05; \*\**P*<0.01; \*\*\**P*<0.001; \*\*\*\**P*<0.0001. *P* values in **d**: +/- vs +/+: *P*=0.2738, -/- vs +/+: *P*=0.8888, -/- vs +/-: *P*=0.1261; *P* values in **e**: +/- vs +/+: *P*=0.0002, -/- vs +/+: *P*=0.0002, -/- vs +/-: *P*=0.5822; *P* values in f: +/- vs +/+: *P*<0.0001, -/- vs +/+: *P*<0.0001, -/- vs +/-: *P*=0.2076.

**Extended Data Fig. 3.**
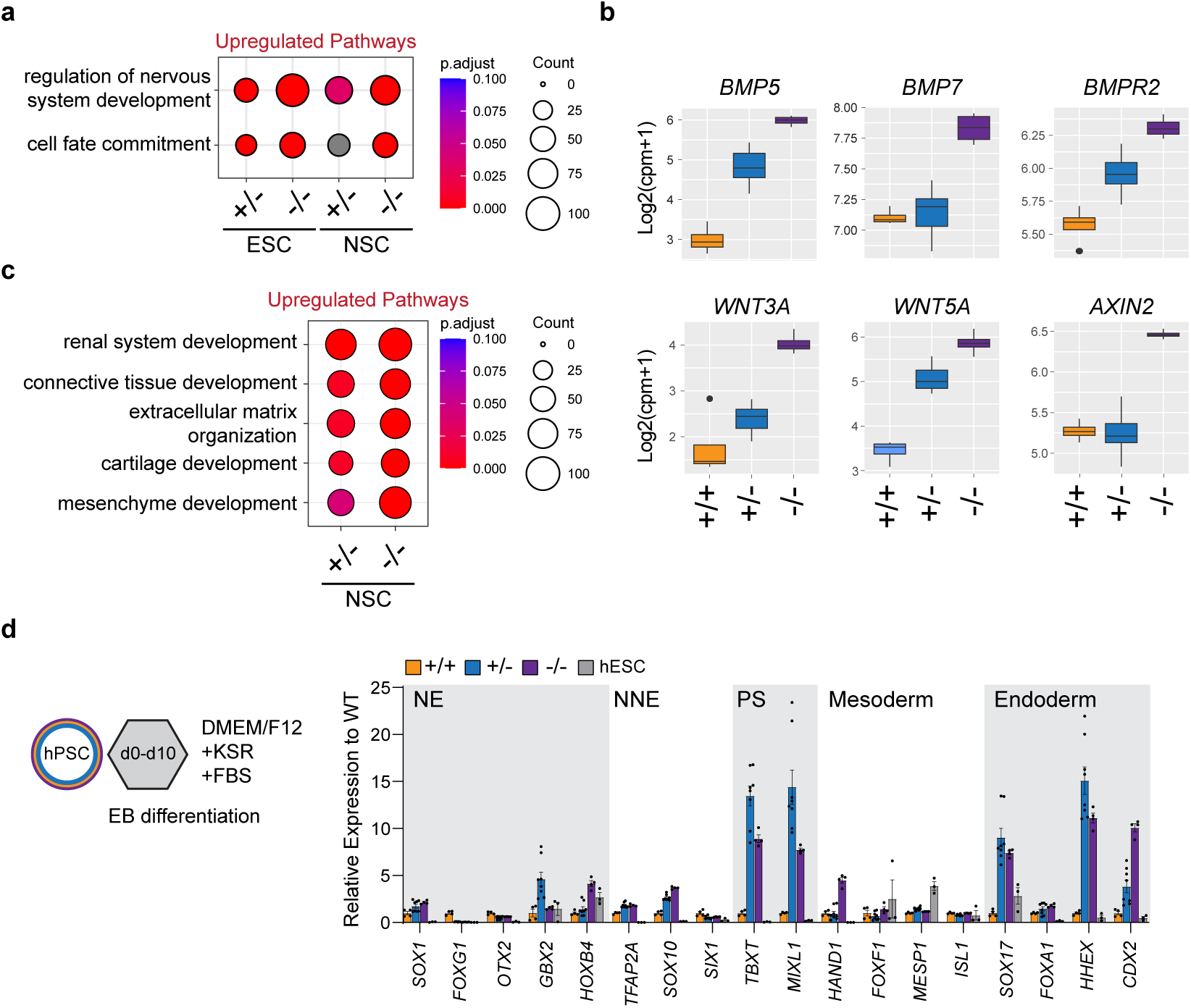
Transcriptomic analysis on WT and ZMYND11 deficient cortical NSCs identified additional pathways related to non-brain development. Related to. Figure 2**. a.** Gene ontology analysis on upregulated genes in ZMYND11 deficient hESCs and NSCs using biological processes. *P*.adjust values were calculated using Benjamini-Hochberg method. **b.** Normalized expression of selected genes in BMP & WNT pathways that are upregulated in ZMYND11 deficient NSCs (*BMP7, BMP5, BMPR2; WNT3A, WNT5A, AXIN2*). **c.** Gene ontology analysis on ZMYND11 deficient NSCs focusing on non-brain development terms. *P*.adjust values were calculated using Benjamini-Hochberg method. **d.** Scheme outlining spontaneous differentiation of WT and ZMYND11 deficient embryoid bodies (left) and qPCR analysis on markers related to different germ layer formation relative to WT (right, mean±SEM, n=4 differentiations for WT and -/-, n=8 differentiations using two clones for +/-). NNE: non-neuroectoderm. NE: neuroectoderm. PS: primitive streak.

**Extended Data Fig. 4.**
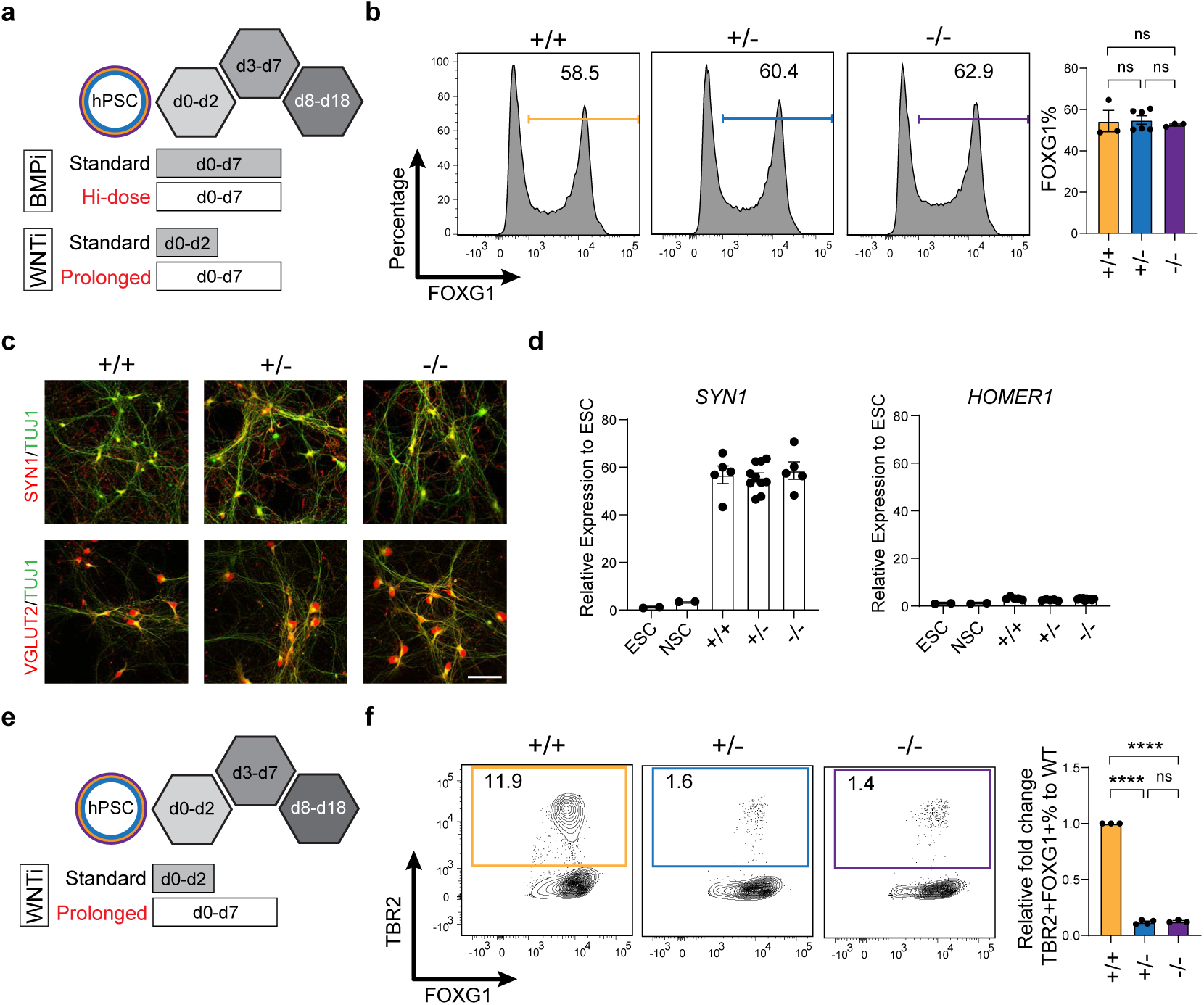
Optimization of differentiation of WT and ZMYND11 deficient cortical NSCs and neurons under high dose BMP and/or WNT inhibition. Related to Figure 2. **a.** Scheme outlining high-dose BMP (LDN193189: 500nM) and WNT inhibition (prolonged XAV939 treatment) strategy during monolayer differentiation. **b.** Intracellular flow cytometry analysis for FOXG1 on WT and ZMYND11 deficient monolayer culture after high-dose BMP and WNT inhibition at d16 (left) with quantification (right, mean±SEM, n=3 differentiations for WT and -/-, n=6 differentiations using two clones for +/-). Y axis shows percentage (“normalized to mode” option in FlowJo). **c.** Immunofluorescence for SYN1, VGLUT2 and TUJ1 on WT and ZMYND11 deficient cortical excitatory neurons after differentiation under high-dose BMP inhibition at d60. Scale bars, 50μm. **d.** qPCR analysis on d60 WT and ZMYND11 deficient cortical excitatory neurons on presynaptic marker *SYN1* (left) and postsynaptic marker *HOMER1* (right) relative to hESCs (mean±SEM, n=2 for hESCs and NSCs, n=5 for WT and -/- neurons, n=10 using two clones for +/- neurons). **e.** Scheme outlining high-dose WNT inhibition only strategy (prolonged XAV939 treatment) during monolayer differentiation. **f.** Intracellular flow cytometry analysis for TBR2 and FOXG1 on WT and ZMYND11 deficient monolayer culture after high-dose WNT inhibition at d16 (left) with quantification relative to WT (right, mean±SEM, n=3 differentiations for WT and -/-, n=4 differentiations using two clones for +/-). Statistics in **d, f**: one-way ANOVA followed by Tukey’s test. ns *P* > 0.05, \**P*<0.05; \*\**P*<0.01; \*\*\**P*<0.001; \*\*\*\**P*<0.0001. *P* values in **d**: +/- vs +/+: *P*=0.9936, -/- vs +/+: *P*=0.9126, -/- vs +/-: *P*=0.8517; *P* values in **f**: +/- vs +/+: *P*<0.0001, -/- vs +/+: *P*<0.0001, -/- vs +/-: *P*=0.7967.

**Extended Data Fig. 5.**
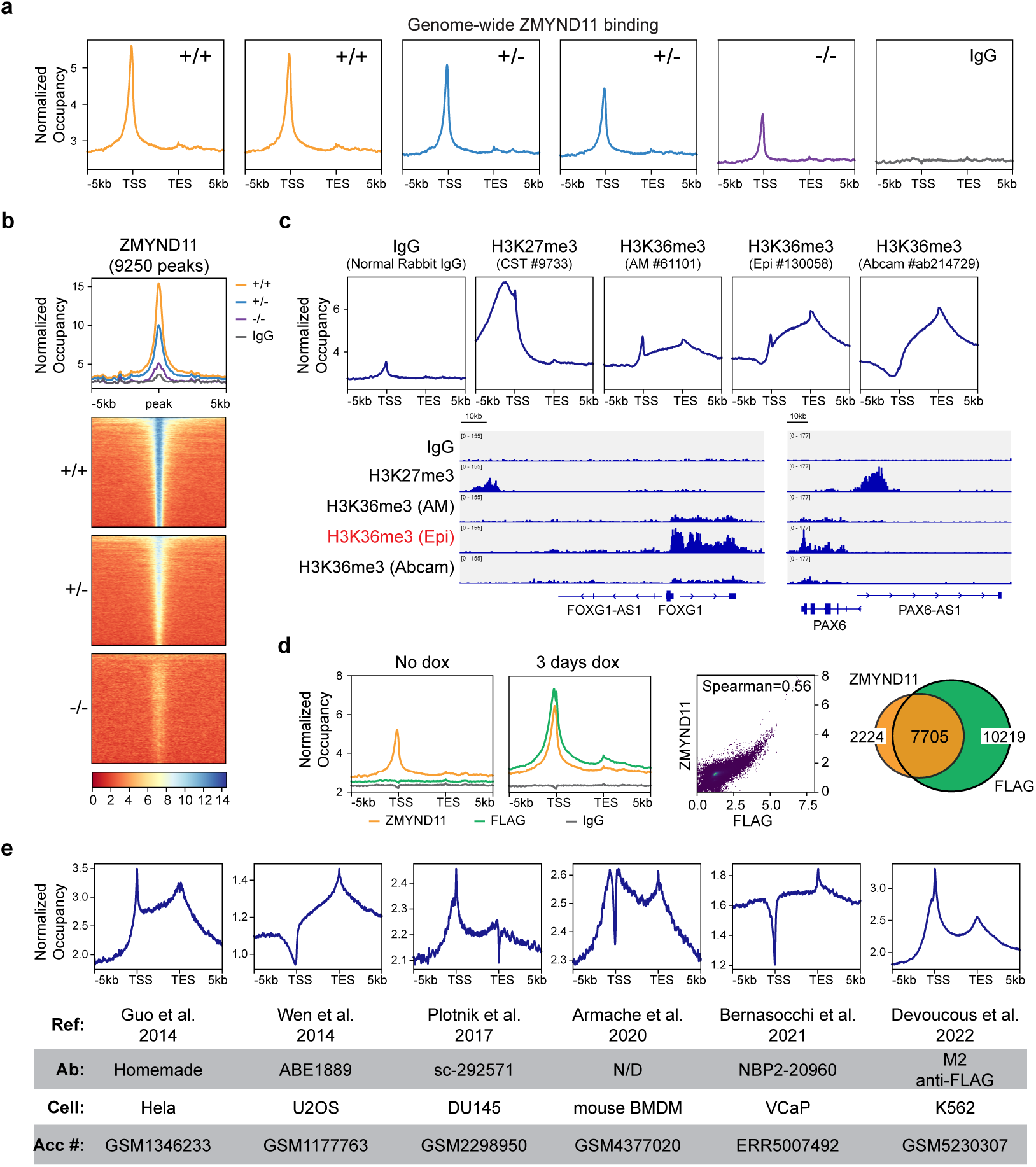
Validation of ZMYND11 binding in WT and ZMYND11 deficient NSCs. Related to Figure 3. **a.** Normalized ZMYND11 binding in WT, ZMYND11 +/- and ZMYND11 -/- cortical NSCs from transcription start sites (TSS) to transcription end sites (TES) ± 5kb (n=2 replicates for WT, n=2 clones for +/-, n=1 clone for -/-). **b.** Normalized ZMYND11-bound peaks and related heatmaps (peaks±5kb). **c.** Normalized H3K36me3 enrichment in WT NSCs using three different antibodies with H3K27me3 enrichment and IgG as controls from transcription start sites (TSS) to transcription end sites (TES) ± 5kb (upper, n=1 for each antibody) and representative tracks (lower, *PAX6, FOXG1*). **d.** Normalized ZMYND11 and FLAG binding in WT NSC with or without dox treatment to overexpress FLAG-ZMYND11 (left), correlation plot of ZMYND11 and FLAG binding after dox treatment (middle, correlation coefficient calculated using Spearman method, n=1 for ZMYND11 without dox, FLAG without dox and ZMYND11 with dox, n=2 replicates for FLAG with dox), and overlap of ZMYND11 and FLAG bound peaks under dox treatment (right). 17924 peaks were called from FLAG binding, and 10626 peaks were called from ZMYND11 binding. **e.** Normalized ZMYND11 binding across publicly available datasets with different cell types.

**Extended Data Fig. 6.**
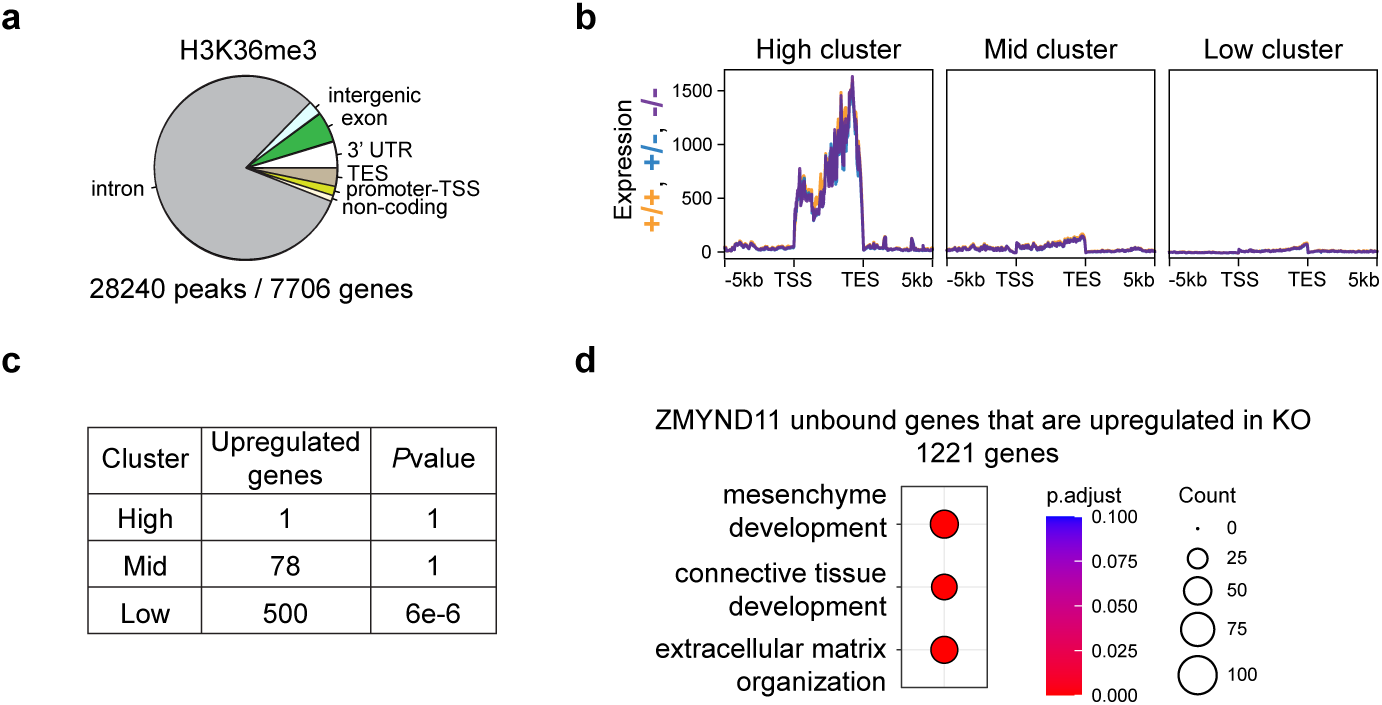
Combined transcriptomic and ZMYND11 binding analysis. Related to Figure 3. **a.** Genomic features distribution, number of called peaks and bound genes of H3K36me3 enrichment in WT NSCs. **b.** Normalized gene expression plot (RPKM) for different ZMYND11 binding clusters (High, Mid, Low). **c.** Table shows the number of upregulated genes in each ZMYND11 binding clusters and hypergeometric *P* values. H: high cluster. M: mid cluster. L: low cluster. **d.** Gene ontology analysis on upregulated genes in ZMYND11 KO that were not bound by ZMYND11 (1221 genes) using biological processes. *P*.adjust values were calculated using Benjamini-Hochberg method.

**Extended Data Fig. 7.**
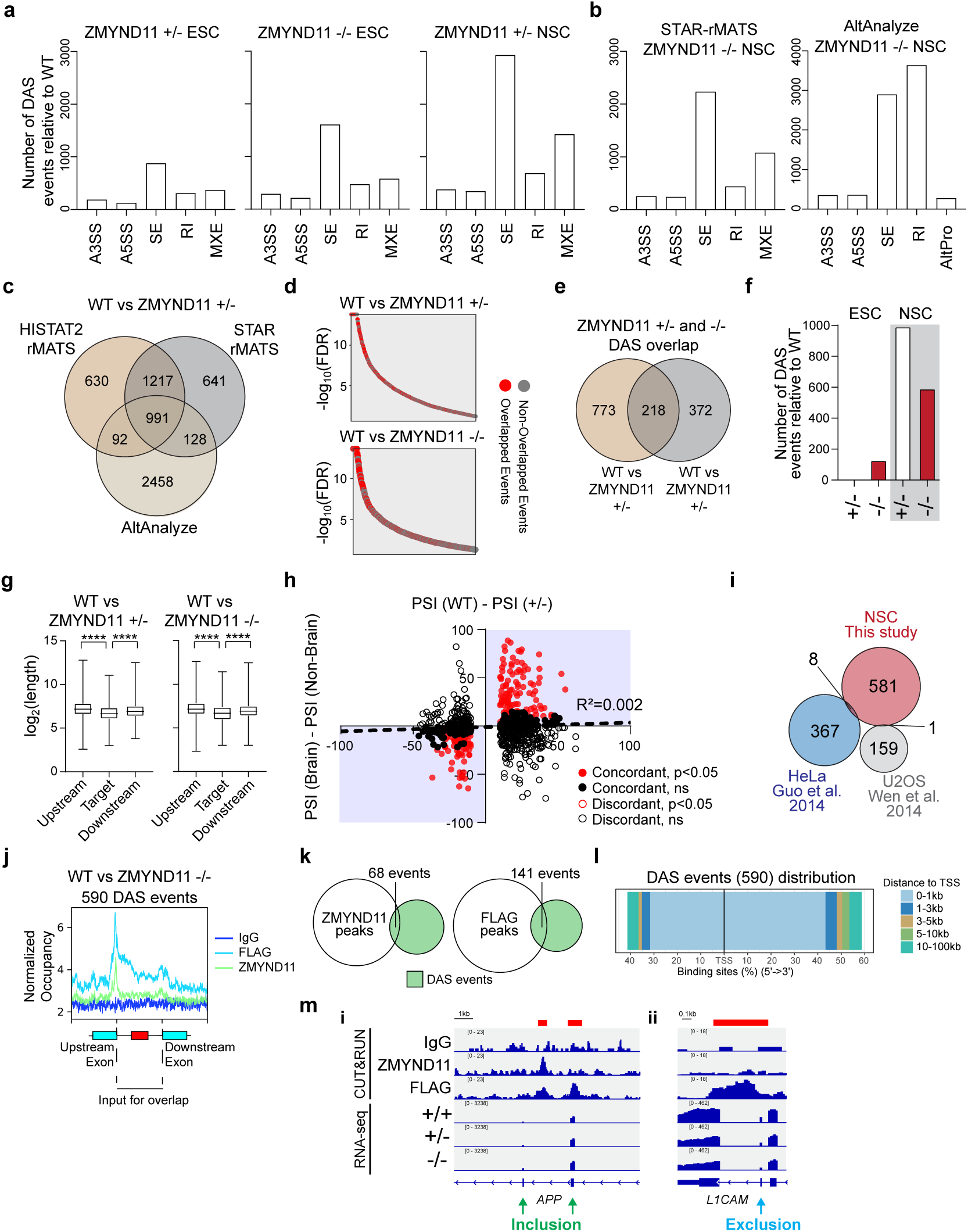
Additional splicing analysis on ZMYND11 deficient cortical NSCs. Related to Figure 4. **a.** Number of significantly changed differential alternative splicing (DAS) events in ZMYND11 +/- ESCs (left), ZMYND11 -/- ESCs (middle) and ZMYND11 +/- NSCs (right) using rMATS prediction and HISAT2 as alignment software. **b.** Number of significantly changed differential alternative splicing (DAS) events in ZMYND11 -/- NSCs using rMATS prediction (left) or AltAnalyze prediction (right) and STAR as alignment software. **c.** Triple overlap of DAS events prediction (HISAT2-rMATS, STAR-rMATS and AltAnalyze) to extract high-confidence DAS events in ZMYND11 +/- NSCs. **d.** Distribution of overlapped events (red) on all HISAT2-rMATS events in ZMYND11 +/- (upper) and ZMYND11 -/- (lower) NSCs. **e.** Overlap of high-confidence DAS events between ZMYND11 +/- NSCs and ZMYND11 -/- NSCs. **f.** Number of high-confidence DAS events across WT and ZMYND11 deficient ESCs and NSCs. **g.** Exon length comparison (log2 transformation) between target exons and upstream or downstream exons in high-confidence DAS events in ZMYND11 +/- (left) and ZMYND11 -/- (right) NSCs. Whiskers show min and max values. **h.** Scatter plot depiction of PSI difference between WT and +/- (x axis) & brain tissues and non-brain tissues (y axis). Similar to Figure 4f. **i.** Overlap of high-confidence DAS events found in this study with DAS events found in previous studies in different cell types. **j**. Normalized ZMYND11 or FLAG-ZMYND11 binding on target exon and its flanking introns. **k**. Overlap between ZMYND11 peaks (left) and FLAG-ZMYND11 peaks (right) with high-confidence DAS events (using genomic coordinates of target exon and its flanking introns). **l**. Distance of high-confidence DAS events to TSS. **m**. Representative tracks of ZMYND11 or FLAG-ZMYND11 binding and related exon expression (i, Green arrow: inclusion events *APP* exon 7 and 8 (left); ii, Blue arrow: exclusion event *L1CAM* exon 28 (right)). Red marks called ZMYND11 or FLAG-ZMYND11 peak. Statistics in **g**: one-way ANOVA followed by Tukey’s test for three groups. \*\*\*\**P*<0.0001. *P* values in **g**: *P*<0.0001.

**Extended Data Fig. 8.**
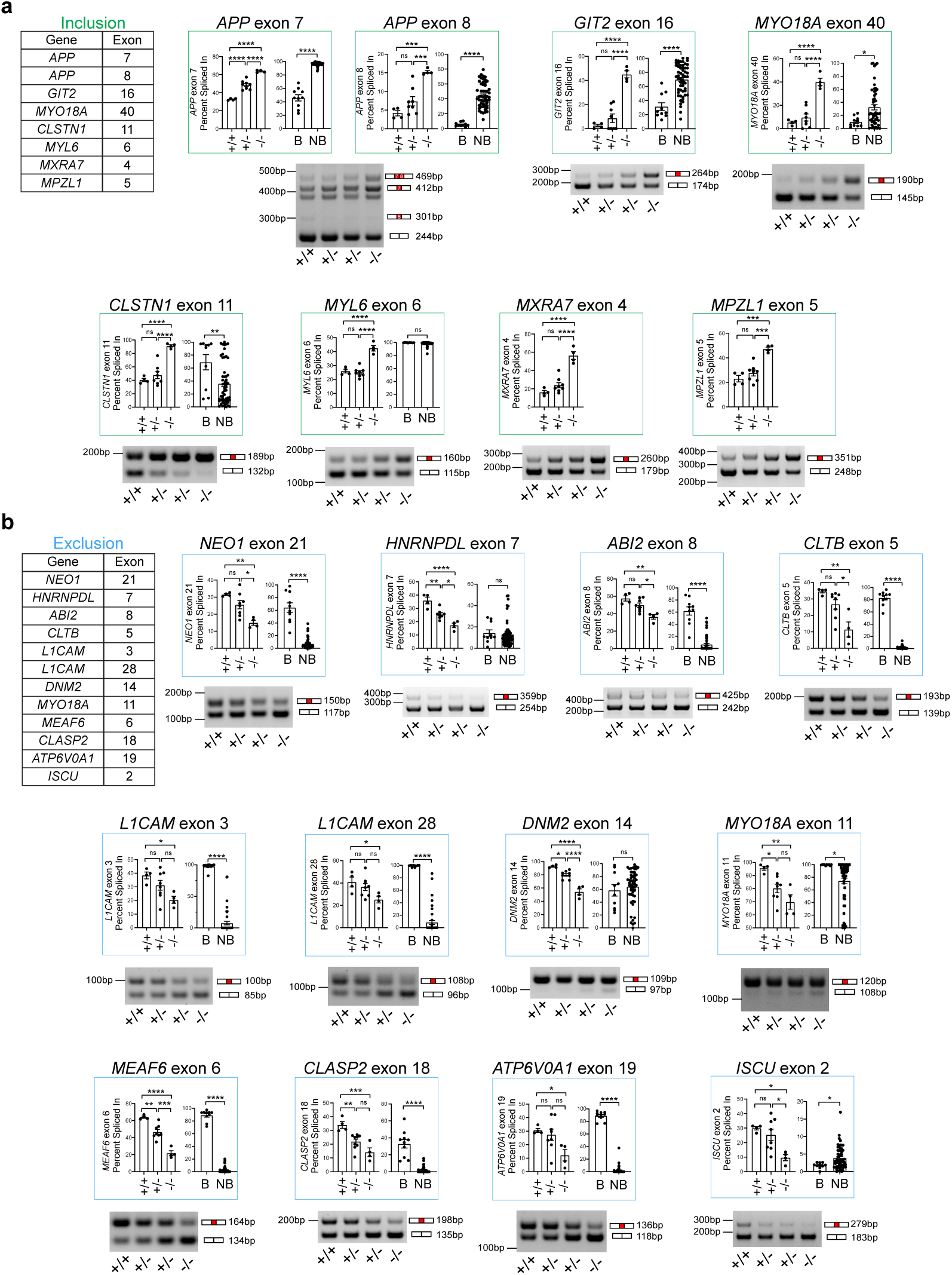
Examples of high-confidence DAS events with PSI changes and validation by semi-quantitative PCR. Related to Figure 4. **a-b.** Example inclusion (**a**) and exclusion (**b**) high-confidence DAS events in ZMYND11 KO with PSI changes (upper) and semi-quantitative PCR validation (lower). B: Brain. NB: Non-Brain. Note that not all events have tissue-specific PSIs in *VastDB* database. Statistics in **a, b**: one-way ANOVA followed by Tukey’s test for three groups, unpaired two-tailed *t* test for two groups. ns *P* > 0.05, \**P*<0.05; \*\**P*<0.01; \*\*\**P*<0.001; \*\*\*\**P*<0.0001. *P* values for *APP* exon 7: +/- vs +/+: *P*<0.0001, -/- vs +/+: *P*<0.0001, -/- vs +/-: *P*<0.0001, NB vs B: *P*<0.0001. *P* values for *APP* exon 8: +/- vs +/+: *P*=0.1634, -/- vs +/+: *P*=0.0002, -/- vs +/-: *P*=0.0009, NB vs B: *P*<0.0001. *P* values for *GIT2* exon 16: +/- vs +/+: *P*=0.4343, -/- vs +/+: *P*<0.0001, -/- vs +/-: *P*<0.0001, NB vs B: *P*<0.0001. *P* values for *MYO18A* exon 40: +/- vs +/+: *P*=0.5970, -/- vs +/+: *P*<0.0001, -/- vs +/-: *P*<0.0001, NB vs B: *P*=0.0222. *P* values for *CLSTN1* exon 11: +/- vs +/+: *P*=0.5687, -/- vs +/+: *P*<0.0001, -/- vs +/-: *P*<0.0001, NB vs B: *P*=0.0049. *P* values for *MYL6* exon 6: +/- vs +/+: *P*=0.8967, -/- vs +/+: *P*<0.0001, -/- vs +/-: *P*<0.0001, NB vs B: *P*=0.1020. *P* values for *MXRA7* exon 4: +/- vs +/+: *P*=0.2327, -/- vs +/+: *P*<0.0001, -/- vs +/-: *P*<0.0001. *P* values for *MPZL1* exon 5: +/- vs +/+: *P*=0.4173, -/- vs +/+: *P*=0.0001, -/- vs +/-: *P*=0.0002. *P* values for *NEO1* exon 21: +/- vs +/+: *P*=0.1892, -/- vs +/+: *P*=0.0017, -/- vs +/-: *P*=0.0146, NB vs B: *P*<0.0001. *P* values for *HNRNPDL* exon 7: +/- vs +/+: *P*=0.0020, -/- vs +/+: *P*<0.0001, -/- vs +/-: *P*=0.0102, NB vs B: *P*=0.7421. *P* values for *ABI2* exon 8: +/- vs +/+: *P*=0.1642, -/- vs +/+: *P*=0.0014, -/- vs +/-: *P*=0.0135, NB vs B: *P*<0.0001. *P* values for *CLTB* exon 5: +/- vs +/+: *P*=0.2324, -/- vs +/+: *P*=0.0019, -/- vs +/-: *P*=0.0134, NB vs B: *P*<0.0001. *P* values for *L1CAM* exon 3: +/- vs +/+: *P*=0.3580, -/- vs +/+: *P*=0.0228, -/- vs +/-: *P*=0.1243, NB vs B: *P*<0.0001. *P* values for *L1CAM* exon 28: +/- vs +/+: *P*=0.6656, -/- vs +/+: *P*=0.0484, -/- vs +/-: *P*=0.1092, NB vs B: *P*<0.0001. *P* values for *DNM2* exon 14: +/- vs +/+: *P*=0.0147, -/- vs +/+: *P*<0.0001, -/- vs +/-: *P*<0.0001, NB vs B: *P*=0.5402. *P* values for *MYO18A* exon 11: +/- vs +/+: *P*=0.0305, -/- vs +/+: *P*=0.0028, -/- vs +/-: *P*=0.1660, NB vs B: *P*=0.0168. *P* values for *MEAF6* exon 6: +/- vs +/+: *P*=0.0027, -/- vs +/+: *P*<0.0001, -/- vs +/-: *P*=0.0001, NB vs B: *P*<0.0001. *P* values for *CLASP2* exon 18: +/- vs +/+: *P*=0.0069, -/- vs +/+: *P*=0.0005, -/- vs +/-: *P*=0.0828, NB vs B: *P*<0.0001. *P* values for *ATP6V0A1* exon 19: +/- vs +/+: *P*=0.8524, -/- vs +/+: *P*=0.0462, -/- vs +/-: *P*=0.0580, NB vs B: *P*<0.0001. *P* values for *ISCU* exon 2: +/- vs +/+: *P*=0.6635, -/- vs +/+: *P*=0.0122, -/- vs +/-: *P*=0.0234, NB vs B: *P*=0.0426.

**Extended Data Fig. 9.**
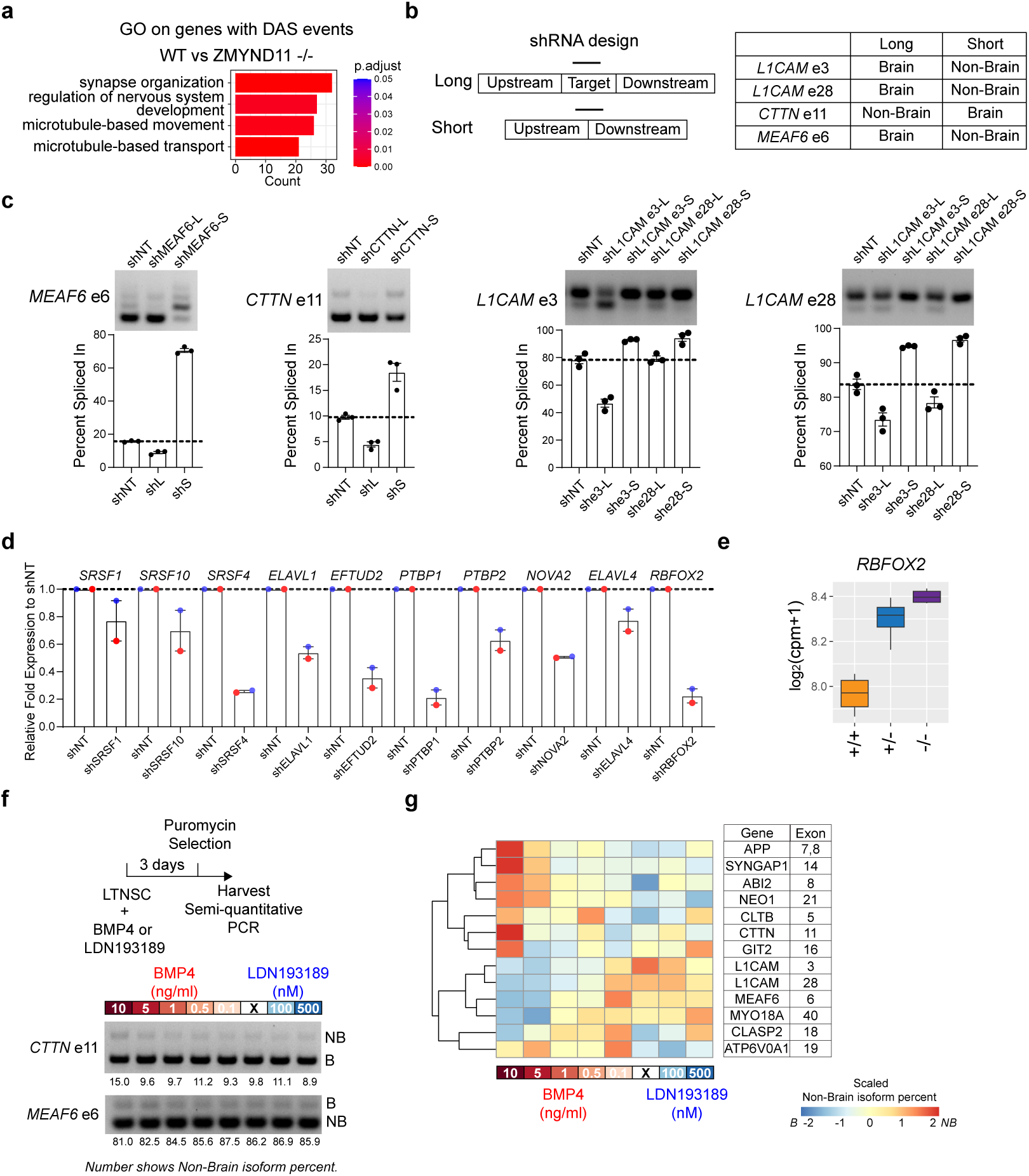
shRNA knockdown experiments to evaluate the cause and the outcome of tissue-specific isoform switch in ZMYND11 KO. **a.** Gene ontology analysis on genes with high-confidence DAS events. *P.*adjust values were calculated using Benjamini-Hochberg method. **b.** Scheme outlining design of isoform specific knockdown shRNA (left) and tissue specificity of isoforms (right). **c.** Semiquantitative PCR for validation of isoform specific knockdown in LTNSCs (upper) and quantification (lower, mean±SEM, n=3 infections). NT: non-targeting. **d.** qPCR for validation of splicing factors knockdown in LTNSCs (n=2 infections, red or blue marks individual batch). **e.** Normalized expression of RBFOX2 in WT and ZMYND11 deficient NSCs from transcriptomic datasets. **f.** Scheme outlining different BMP4 and BMP inhibitor treatment on LTNSCs that were harvested for semiquantitative PCR to investigate the impacts of BMP signaling on tissue-specific splicing pattern (upper), and semi-quantitative PCR gels on *CTTN* exon 11 and *MEAF6* exon 6 (lower). **g.** Heatmap for **f** using non-brain isoform percent values (row-wise scaled by Z-score transformation) on selected high-confidence DAS events that showed match between non-brain tissues and ZMYND11 KO.

**Extended Data Fig. 10.**
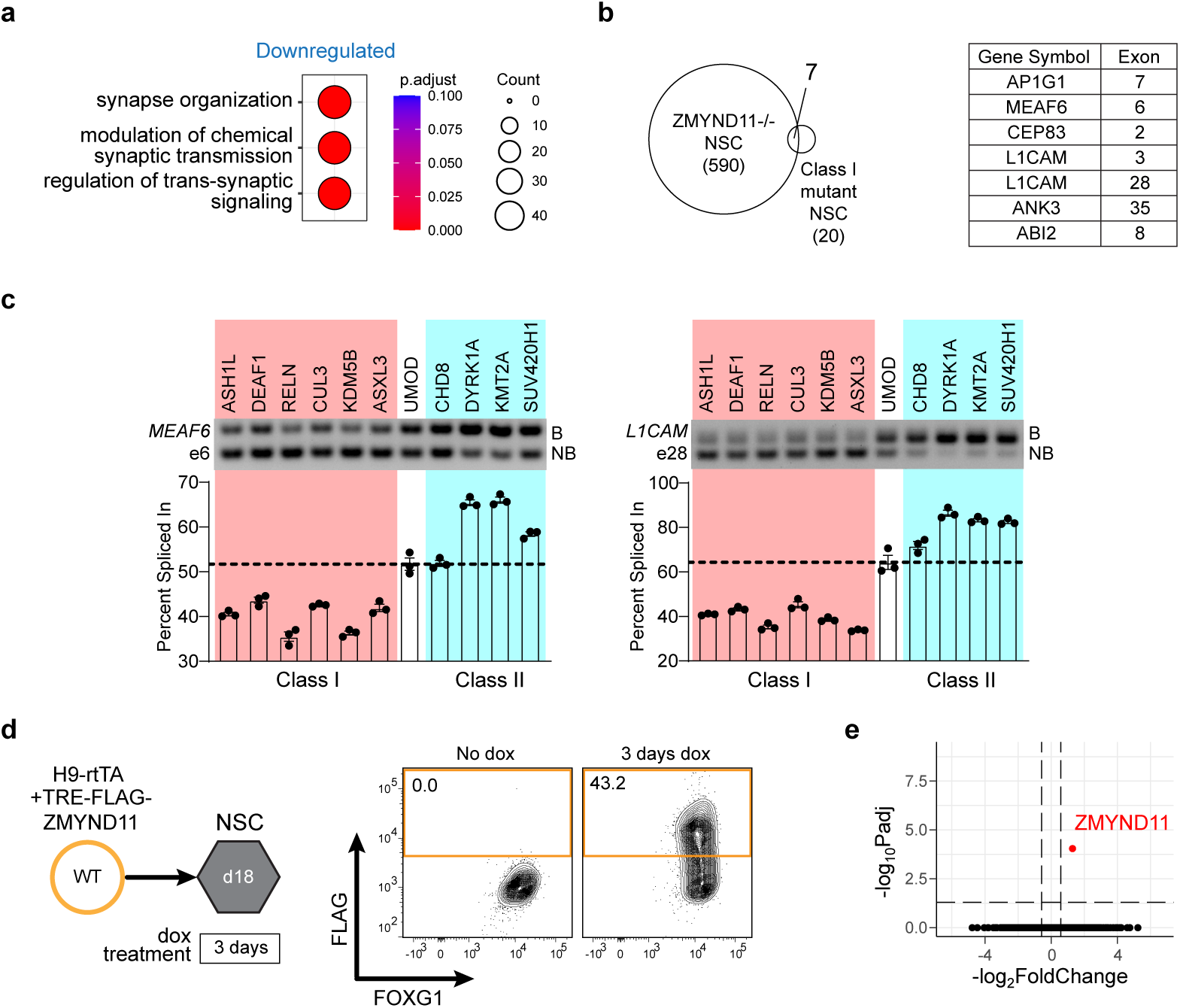
Further characterization of Class I mutant differentiations. **a.** Gene ontology analysis on downregulated genes in Class I mutant cortical NSCs using biological processes. *P*.adjust values were calculated using Benjamini-Hochberg method. **b.** Overlap of high-confidence DAS events between Class I mutant NSCs and ZMYND11 -/- NSCs (left) and table of overlapped events (right). **c.** Semiquantitative PCR of *MEAF6* exon 6 (left) and *L1CAM* exon 28 (right) on both Class I mutant and Class II mutant NSCs and quantification (mean±SEM, n=3 differentiations). **d.** RNA-seq on WT differentiation that overexpressed FLAG-ZMYND11 for 3 days (left) and quantification of FLAG+ population by intracellular flow cytometry (right). **e.** Volcano plot shows only ZMYND11 was differentially expressed when overexpressing FLAG-ZMYND11 for 3 days in WT cortical NSCs.

