## Supplementary material for "ZMYND11 Functions in Bimodal Regulation of Latent Genes and Brain-like Splicing to Safeguard Corticogenesis": Chang_et_al_2024_ZM11_Description_of_Supplementary_Information_241014.docx

**Supplementary Table 1:** Differential gene expression analysis on ZMYND11 mutant hESCs and NSCs, and Class I mutant NSCs.

**Supplementary Table 2:** ZMYND11 bound and H3K36me3 enriched peaks and genes on WT NSCs, and FLAG bound and ZMYND11 bound peaks on WT NSCs overexpressing FLAG-ZMYND11.

**Supplementary Table 3:** High-confidence differential alternative splicing (DAS) events on ZMYND11 mutant hESCs and NSCs, and Class I mutant NSCs.

**Supplementary Table 4:** All primer, shRNA and sgRNA sequences and Class I & II mutant clone information.

**Supplementary Table 5:** Antibodies and dilution used in this study.
